# *APC* and *P53* mutations synergize to create a therapeutic vulnerability to NOTUM inhibition in advanced colorectal cancer

**DOI:** 10.1101/2022.07.26.501557

**Authors:** Yuhua Tian, Xin Wang, Zvi Cramer, Joshua Rhoades, Katrina N. Estep, Xianghui Ma, Stephanie Adams Tzivelekidis, Bryson W Katona, F Brad Johnson, Zhengquan Yu, Mario Andres Blanco, Christopher Lengner, Ning Li

**Affiliations:** Department of Biomedical Sciences, School of Veterinary Medicine, University of Pennsylvania, Philadelphia, PA, United States; Department of Pathology and Laboratory Medicine, Perelman School of Medicine, University of Pennsylvania, Philadelphia, PA, United States; Children’s Hospital of Philadelphia; Department of Medicine, Perelman School of Medicine, University of Pennsylvania, Philadelphia, PA, United States; China Agricultural University; University of Pennsylvania School of Veterinary Medicine; University of Pennsylvania

## Abstract

Colorectal cancer (CRC) is a leading cause of cancer-related deaths globally, with the majority of cases initiated by inactivation of the APC tumor suppressor. This results in the constitutive transcriptional activation of the canonical WNT signal transduction pathway effector β-Catenin, along with induction of WNT feedback inhibitors, including the extracellular palmitoleoyl-protein carboxylesterase NOTUM. Here, we show that NOTUM retains cell-autonomous tumor suppressive activity in APC-null adenomatous lesions despite constitutive β-Catenin activation. Strikingly, we find that NOTUM becomes an obligate oncogene upon subsequent P53 inactivation during the adenoma-adenocarcinoma transition, and that these phenotypes are WNT-independent, resulting from differential activity of NOTUM upon its enzymatic targets Glypican 1 and 4 in early vs. late-stage disease, respectively. Ultimately, preclinical mouse models of CRC and human tumoroid cultures demonstrate that pharmacological inhibition of NOTUM is highly effective in arresting primary adenocarcinoma growth and inhibiting metastatic colonization of distal organs. The finding that a single agent targeting an extracellular enzyme is effective in treating highly aggressive tumors make NOTUM a novel therapeutic vulnerability in advanced colorectal adenocarcinomas.

## Introduction

Advanced colorectal cancer (CRC) represents a major unmet medical need as the third most commonly diagnosed cancer in the US, and the third leading cause of cancer-related deaths, with 5-year survival rates around 10%. The majority (>80%) of CRC is driven by loss-of-function mutations in the tumor suppressor APC ^[1–3]^. APC inactivation results in the constitutive transcriptional activation of the canonical WNT signal transduction pathway effector β-CATENIN ^[4, 5]^. β-CATENIN, in turn, induces expression of genes that drive stem cell cycling and self-renewal including *MYC*, *LGR5*, and *CCND1* ^[6]^. However, APC loss also activates feedback inhibitors of canonical WNT signaling, including the extracellular palmitoleoyl-protein carboxylesterase NOTUM, known to inhibit WNT signaling at the ligand/receptor level in normal intestinal epithelium. NOTUM inhibition of the canonical WNT pathway has been proposed to occur through two distinct mechanisms. First, studies in *Drosophila* revealed that Notum can enzymatically cleave two glypicans, Dally, and Dally-like protein (Dlp) from the cell surface ^[7–9]^. These glypicans potentiate WNT receptor-ligand interactions, and thus Notum antagonizes canonical WNT pathway activity in this model ^[7–9]^. NOTUM cleavage of mammalian glypicans (which exist as a family of six paralogous orthologs to *Drosophila* glypicans) was subsequently confirmed ^[10]^. A second model for NOTUM antagonism of WNT signaling, which is not mutually exclusive from the first, is through NOTUM’s ability to directly cleave the palmitoleate tails from WNT ligands, thus rendering them incapable of productive interaction with their Fzd receptors ^[11]^.

Both of these models for NOTUM function posit that its ability to antagonize WNT activity occurs at the level of receptor-ligand interaction at the cell surface. Therefore, inactivation of APC, which leads to constitutive β-CATENIN activation due to inactivation of the destruction complex downstream of receptor-ligand interactions, is predicted to render APC-null cells insensitive to the extracellular activity of NOTUM upon its WNT ligand or glypican cleavage targets. Consistent with this, recent studies investigating the role of NOTUM in APC-null intestinal stem cell clones in mice suggest that its ability to attenuate canonical WNT activity confers a competitive advantage to APC-null cells over their wildtype neighbors ^[12, 13]^. This model posits that NOTUM produced by a clone that has inactivated APC (and thus should no longer depend on WNT receptor-ligand interactions for β-CATENIN transcriptional activity) can act upon neighboring stem cells that remain reliant on WNT ligands for their proliferative self-renewal in order to suppress their growth, thus favoring outgrowth of the APC-null clone to initiate adenoma formation ^[13]^.

Here, we evaluated the functional contribution of NOTUM in genetically-defined tumor organoid (tumoroid) models of CRC and made two striking observations. First, NOTUM retains potent, cell-autonomous tumor suppressive activity in APC-null epithelial cells, despite constitutive downstream activation of β-CATENIN. Second, upon subsequent inactivation of the P53 tumor suppressor during the adenoma-to-adenocarcinoma transition, NOTUM activity paradoxically becomes highly oncogenic, despite being tumor suppressive in the presence of inactivating *Apc* or *Trp53* mutations alone. Consistently, we find that as CRC progresses and acquires subsequent driver mutations (e.g., oncogenic *Kras* activation and *Smad4* inactivation) these cancers remain dependent on oncogenic NOTUM activity.

We addressed the molecular mechanism whereby NOTUM mediates its oncogenic effects in advanced CRC, as established models for NOTUM inhibition of canonical WNT pathway activity cannot account for this phenomenon. We found that differential activity of Glypicans GPC1 and GPC4 in APC-null adenomas and advanced colorectal adenocarcinomas accounted for the tumor suppressor-to-oncogene switch of NOTUM activity. Ultimately, preclinical mouse models of invasive and metastatic CRC demonstrated that pharmacological inhibition of NOTUM is highly effective as a single agent in arresting primary adenocarcinoma growth and inhibiting metastatic colonization of distal organs. Analysis of human CRC metadata and tumoroid cultures support our findings in mouse models and provide the impetus for evaluating NOTUM inhibition as a genotype-specific therapeutic intervention in advanced colorectal cancers.

## Results

### *Notum* is an obligate oncogene in colorectal adenocarcinoma

The majority of colorectal cancers exhibit inactivation of the APC tumor suppressor, and APC reactivation in established tumors is sufficient to arrest tumor growth, restore differentiation and re-establish normal tissue architecture, even in the presence of oncogenic *Trp53* and *Kras* driver mutations ^[14]^. We therefore reasoned that the immediate effects of APC inactivation may represent unexplored therapeutic vulnerabilities in later stages of advanced colorectal adenocarcinoma. To this end we performed transcriptome profiling of epithelial colon organoids immediately after APC inactivation and cross-referenced gene expression changes to the transcriptomes of several mouse models of colon cancer, including colon tumors in an inflammation-induced CRC model (AOM/DSS), a model of familial adenomatous polyposis (FAP, the *Apc^Min/+^* mouse), and an implantation-based model of metastatic adenocarcinoma using *Apc/Trp53/Kras^G12D^/Smad4* quadruple-mutant tumor organoids^[15]^(*APKS* tumoroids) (**Fig. 1a**). Of the roughly 150 genes whose expression is acutely induced upon APC loss and remains elevated in these colon cancer models (**Supplemental Table 1**), the palmitoleoyl-protein carboxylesterase *Notum* stood out as it has previously been implicated in initiation of adenoma formation^[13]^. Further, as an extracellular enzyme it represents a potentially tractable therapeutic target.

**Fig. 1.**
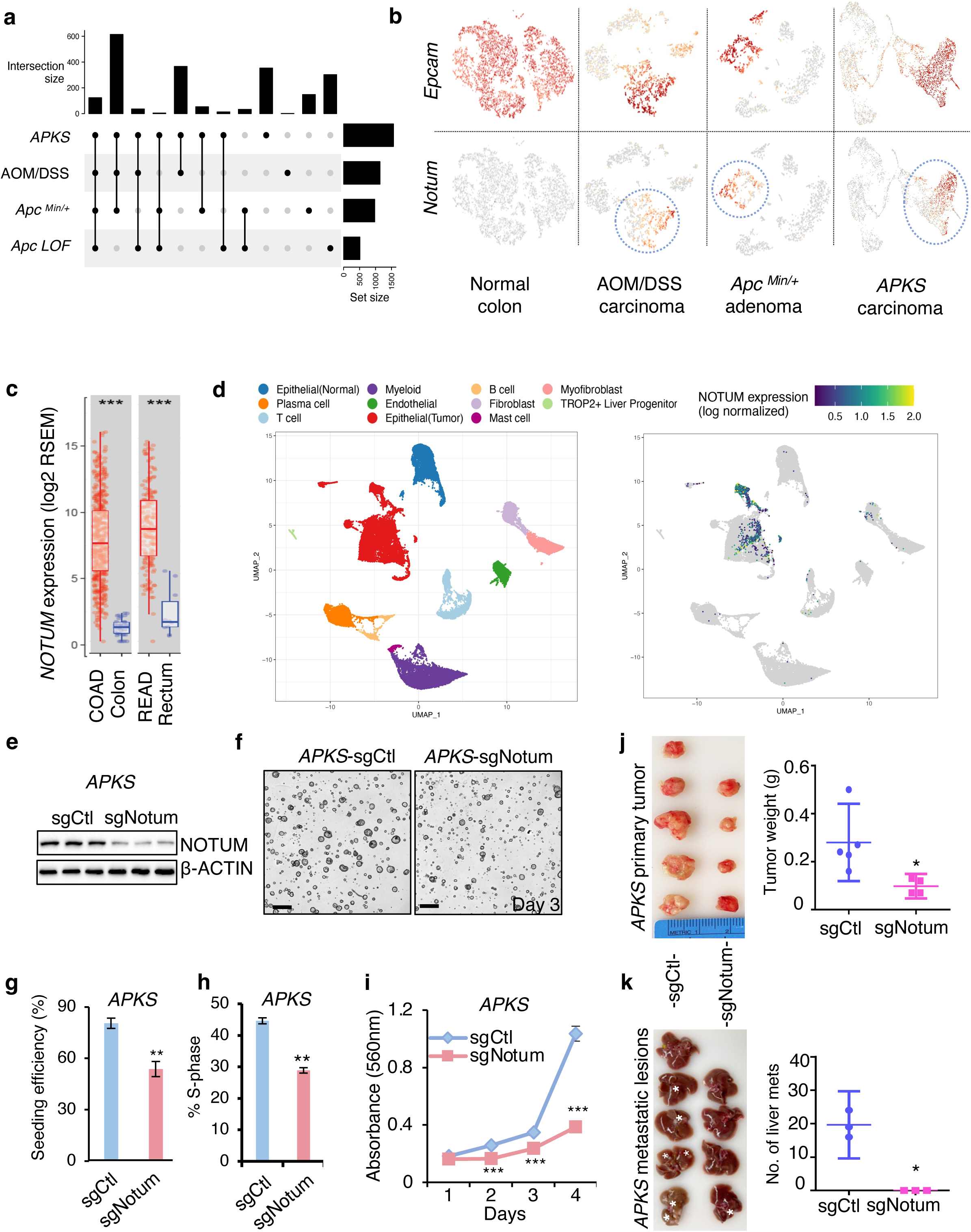
*Notum* is a cell-autonomous oncogene in colorectal adenocarcinoma. **(a)** UpSet plot showing common differentially expressed genes (p-value < 0.05, log2FC > 1) in tumor cells from mouse models of CRC relative to normal colonic epithelium. *APKS*: adenocarcinoma resulting from orthotopic implantation of *Apc/Trp53/Kras/Smad4* quadruple-mutant tumoroids into syngeneic mouse colon. AOM/DSS: inflammation-induced colon adenocarcinoma. *Apc^Min/+^*: colon adenoma induced by spontaneous loss of *Apc* heterozygosity in the *Apc^min/+^* mouse model. *Apc LOF*: colonic epithelium after acute loss-of-function of *Apc* in an *Apc^flox/flox^::Villin-CreER* mouse 5 days after tamoxifen-induced *Apc* inactivation. **(b)** Uniform manifold approximation projections (UMAP) of single cell RNA sequencing (scRNAseq) datasets from tumor models in (**a**), with epithelial/carcinoma cells expressing *Epcam* (top). *Notum* expression (bottom) is restricted to epithelial/carcinoma cells (circled). **(c)** *NOTUM* expression in the Cancer Genome Atlas (TCGA) colon (COAD) and rectal (READ) adenocarcinoma datasets. **(d)** Left: UMAP representation the cell populations of human CRC primary tumor and adjacent normal epithelium. Cells are colored according to their cell types. (n=17). Right: UMAP showing *NOTUM* expression restricted to human carcinoma cells. **(e)** Western blot analysis for NOTUM in *APKS* tumoroids infected with control sgRNA (sgCtl) or *Notum* sgRNA(sgNotum). β-ACTIN was used as a loading control. **(f)** Images of *APKS* tumoroids from (**e**) three days after seeding (n=3). Scale bar: 200 *μ*m. **(g)** Clonal seeding efficiency of single *APKS* cells from (**e/f**). **(h)** Flow cytometric analysis of EdU incorporation in *APKS* tumoroids infected with control sgRNA (sgCtl) or *Notum* sgRNA (sgNotum) (n=3). **(i)** MTT proliferation assays of *APKS* tumoroids (n=3). **(j)** Left: Primary tumors formed 8 weeks after orthotopic injection of *APKS* tumoroids into the distal colon of syngeneic recipient mice (n=5 in sgCtl group, n=4 in sgNotum group). Right: Weight of primary tumors from the left. **(k)** Left: Livers of the mice bearing primary *APKS* tumors from (**j**). White asterisks show metastatic liver lesions. Right: The number of macrometastases visible in the liver of mice from (**j**). For all panels: \**P*<0.05, \*\**P*<0.01, \*\*\**P*<0.001, Student’s *t*-test.

High NOTUM expression was confirmed in mouse tumors, where its expression is limited to epithelial/tumor cells (**Fig. 1b** and **Extended Data Fig. 1a, b**). We also confirmed strong activation of NOTUM in response to acute loss of APC or activation of the WNT effector β-CATENIN in the colonic epithelium of mice, consistent with reports that *Notum* is a direct β-CATENIN target gene ^[16]^ (**Extended Data Fig. 1c**). Consistent with the mouse data, *NOTUM* expression is significantly elevated in colon and rectal cancers in the human Cancer Genome Atlas data ^[3]^ (TCGA COAD and READ, respectively) relative to normal tissue (**Fig. 1c**). We also examined single cell transcriptome profiles of primary human colon adenocarcinomas and confirmed that *NOTUM* expression is largely restricted to carcinoma cells and almost entirely absent in cells of the tumor microenvironment (TME) (**Fig. 1d**). Finally, examining a group of paired human primary tumors and their cognate liver metastases revealed further increases in *NOTUM* expression in the metastatic lesions (**Extended Data Fig. 1d**). These findings demonstrate that *NOTUM* expression is induced in human and murine colonic epithelium in response to APC inactivation, and remains elevated throughout the transition from adenoma to invasive adenocarcinoma in both human disease and mouse models thereof.

We initially evaluated the carcinoma cell-autonomous function of NOTUM in a model of invasive and metastatically competent colon adenocarcinoma: *APKS* tumoroids generated by engineering oncogenic inactivating mutations in *Apc*, *Trp53*, and *Smad4* using CRISPR/Cas9 ^[15]^, and a targeted, conditional activating mutation in *Kras* (*Kras^G12D^*) ^[17]^. Upon introduction of gRNAs targeting *Notum*, bulk tumoroid cultures grew slower and had significantly diminished clonal tumoroid seeding efficiencies, a proxy for tumor-initiating capacity (**Fig. 1e-i, Extended Data Fig. 1e, f**). Conversely, gain-of-function (GOF) experiments in *Apc/Trp53* (*AP*), *Apc/Trp53/Kras* (*APK*), and *APKS* tumoroids revealed that NOTUM overexpression is able to accelerate proliferation across this allelic series (**Extended Data Fig. 1g, h**). To test whether NOTUM has similar oncogenic activity *in vivo*, we performed both subcutaneous and orthotopic tumor formation assays with *APKS* tumoroids (the latter using endoscope-guided orthotopic implantation of tumoroids into the colonic mucosa of syngeneic mice^[18]^) and NOTUM loss-of-function (LOF). These assays revealed significant decreases in primary tumor growth, and, in the orthotopic model, a complete block of macrometastases forming in the liver as a result of NOTUM LOF (**Fig. 1j, k** and **Extended Data Fig. 1i-m**). These experiments were performed with bulk tumoroid cultures receiving gRNAs targeting *Notum*. Attempts at establishing subclonal cultures with biallelic *Notum* inactivation failed. One hypomorphic subclonal line exhibiting an approximately 80% reduction in NOTUM levels was stabilized (referred to as *Notum APKS^Hypomorph^*). This line grew poorly *in vitro* and completely failed to generate tumors *in vivo* (**Extended Data Fig. 1n, o**). Taken together, these data suggest that *Notum* may be an obligate oncogene in colorectal adenocarcinoma.

### NOTUM maintains tumor suppressive activity upon APC inactivation

The oncogenic activity of NOTUM in these tumoroid models was surprising and counterintuitive given that NOTUM is an established negative regulator of canonical WNT signaling in the intestinal epithelium ^[11, 13]^, and WNT hyperactivation is a primary driver of colon cancer. We thus returned to re-evaluate NOTUM function in normal organoids and early adenomatous tumoroids upon APC inactivation. NOTUM LOF in bulk wildtype colon organoid culture via CRISPR/Cas9 resulted dramatic increases in clonal organoid seeding efficiency, organoid growth, and canonical WNT/β-Catenin intestinal stem cell target gene expression (**Fig. 2a-c** and **Extended Data Fig. 2a**). Conversely, NOTUM overexpression potently inhibited proliferation and WNT target gene expression (**Fig. 2d, e** and **Extended Data Fig. 2b, c**). These findings are consistent with the literature indicating that Notum negatively regulates the canonical WNT pathway in the normal intestinal stem cell compartment ^[11, 19]^.

**Fig. 2.**
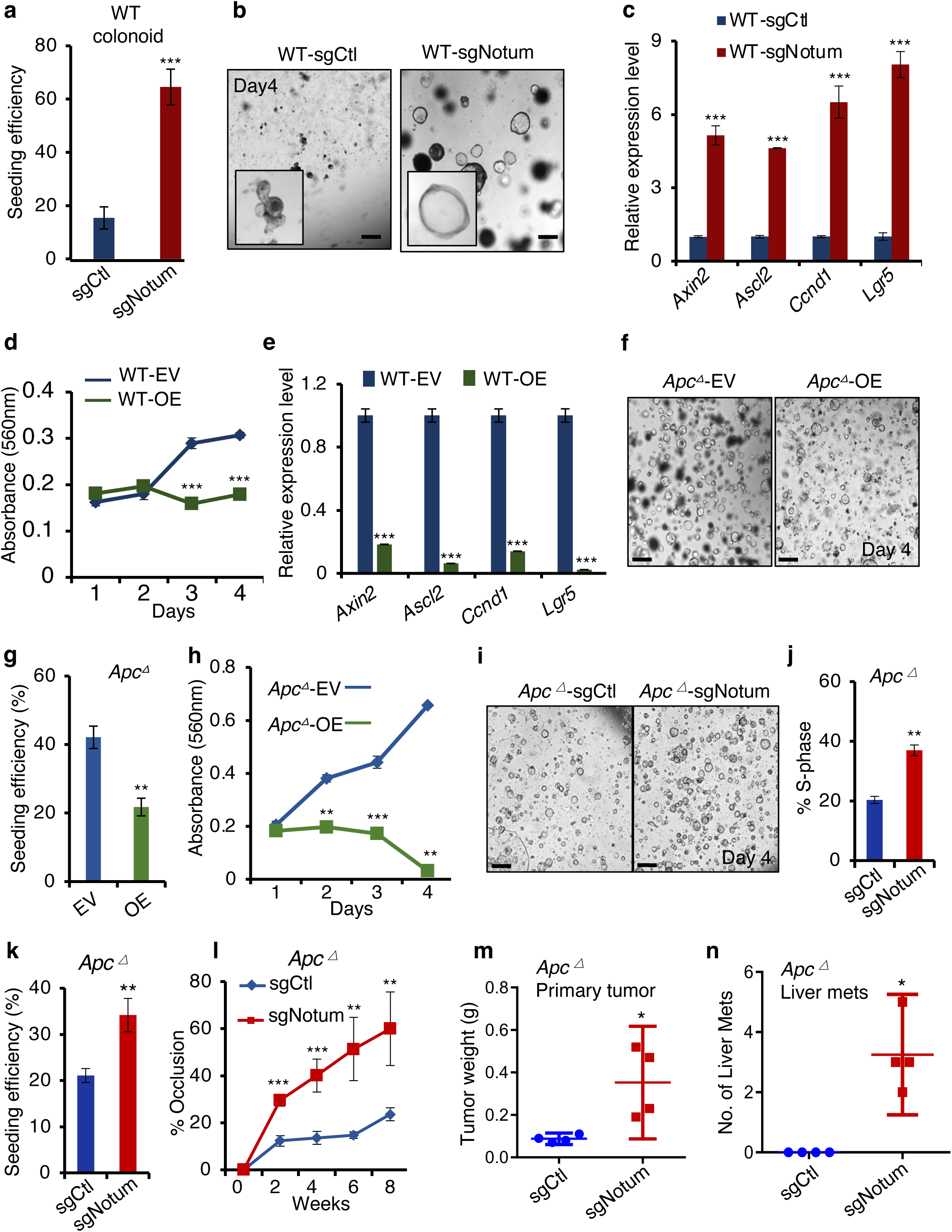
NOTUM has potent tumor suppressive activity in normal and APC-mutant colon tumoroids. **(a)** Clonal seeding efficiency of wildtype (WT) colon organoids infected with control sgRNA (sgCtl) or *Notum* sgRNA (sgNotum) (n=3). **(b)** Images of WT organoids infected with control sgRNA (sgCtl) or *Notum* sgRNA (sgNotum) four days after seeding (n=3). Scale bar: 100 *μ*m. **(c)** Quantitative RT-PCR analysis of canonical Wnt/ β-CATENIN pathway target genes including *Axin2, Ascl2, Ccnd1,* and *Lgr5* in organoids from (**b**). **(d)** MTT proliferation assays of WT organoids infected with empty lentiviral vector (EV) or *Notum* overexpression vector (OE) (n=3). **(e)** Relative expression level of Wnt pathway target genes including *Axin2, Ascl2, Ccnd1* and *Lgr5* in WT organoids infected with empty vector (EV) or *Notum* overexpression plasmid (OE) (n=3). **(f)** Images of *Apc* mutant tumoroids infected with empty vector (EV) or vector overexpressing *Notum* four days after seeding (n=3). Scale bar: 200 *μ*m. **(g)** Seeding efficiency of *Apc* mutant tumoroids from (**f**). **(h)** MTT proliferation assays of *Apc* mutant tumoroids from (**f**) (n=3). **(i)** Images of *Apc* mutant tumoroid cultures infected with control sgRNA (sgCtl) or *Notum* sgRNA (sgNotum) four days after seeding (n=3). Scale bar: 200 *μ*m. **(j)** The percentage of EdU+ cells in cultures from (**i**). **(k)** Clonal seeding efficiency of cells from tumoroids in (**i**). **(l)** Percent lumen occlusion was measured after orthotopic implantation of *Apc-*mutant tumoroids infected with control sgRNA (sgCtl) or *Notum* sgRNA (sgNotum) (n=4). **(m)** Weight of primary, orthotopic *Apc* mutant colon tumors with or without *Notum* loss of function (n=4). **(n)** The number of macroscopically observable liver metastases in mice harboring orthotopic from (**l, m**). For all panels: \**P*<0.05, \*\**P*<0.01, \*\*\**P*<0.001, Student’s *t*-test.

We performed analogous experiments in adenomatous tumoroids with inactivating *Apc* mutations introduced by CRISPR/Cas9 (*Apc*^△^). Remarkably, NOTUM gain-of-function in *Apc*^△^ tumoroids resulted in decreased clonal tumoroid seeding efficiency, proliferation, and Wnt target gene expression while NOTUM loss-of-function had the converse effect (**Fig. 2f-k** and **Extended Data Fig. 2d, e**). We confirmed these findings in a second model of *Apc* LOF: tumoroids isolated from *Apc^flox/flox^::Villin-CreER* mice after Cre activation (**Extended Data Fig. 2f, g**). Given that APC loss stabilizes β-Catenin resulting in its constitutive transcriptional activation, these findings are unexpected, as NOTUM-mediated Wnt antagonism is expected to occur extracellularly at the level of receptor-ligand interaction (and thus *Apc*^△^ cells might be expected to be insensitive to NOTUM activity) ^[13]^. Recent studies suggest that NOTUM induction in an *Apc*^△^ intestinal stem cell confers a competitive advantage over surrounding wildtype epithelial cells via juxtacrine/paracrine activity ^[12, 13]^. Therefore, we next asked how NOTUM loss influences tumorigenesis of *Apc*^△^ tumoroids *in vivo*. Endoscope-guided orthotopic implantation of *Apc*^△^ tumoroids with or without NOTUM LOF revealed that NOTUM retains potent tumor suppressive activity in the absence of APC function. As expected, *Apc*^△^ tumoroids form only small adenomas in the colonic epithelium. In contrast, loss of NOTUM not only results in massive increases in primary tumor size, it endows these tumors with metastatic capacity (**Fig. 2l-n** and **Extended Data Fig. 2h-l**). Thus, NOTUM retains potent tumor suppressive activity in adenomatous lesions driven by APC inactivation despite constitutive β-Catenin activation.

### Novel tumor suppressive and oncogenic functions of NOTUM require extracellular enzymatic activity

Because our findings in *Apc*^△^ tumoroids, and more aggressive *AP/APK/APKS* tumoroids were unexpected based on known models of NOTUM antagonizing canonical Wnt signaling via its extracellular enzymatic activity, we next tested whether NOTUM’s tumor suppressive activity in *Apc*^△^ tumoroids and oncogenic activity in *AP/APK/APKS* tumoroids require enzymatic activity. To this end, we overexpressed wildtype or catalytically dead (S239A^[11]^) NOTUM in *Apc*^△^ tumoroids and found that NOTUM S239A was unable to suppress proliferation and clonal seeding efficiency (**Fig.3 a-c** and **Extended Data Fig. 3a**). Conversely, we overexpressed wildtype and S239A NOTUM in the hypomorphic APKS subclone (with reduced endogenous NOTUM levels, **Extended Data Fig. 1o**), and found that, while wildtype NOTUM increased tumoroid seeding efficiency and proliferation, the catalytically dead NOTUM did not (**Fig. 3d-f** and **Extended Data Fig. 3b**). Thus, catalytic activity is required for both NOTUM’s tumor suppressive activity in *Apc*^△^ tumoroids and for oncogenic activity in *APKS* tumoroids.

**Fig. 3.**
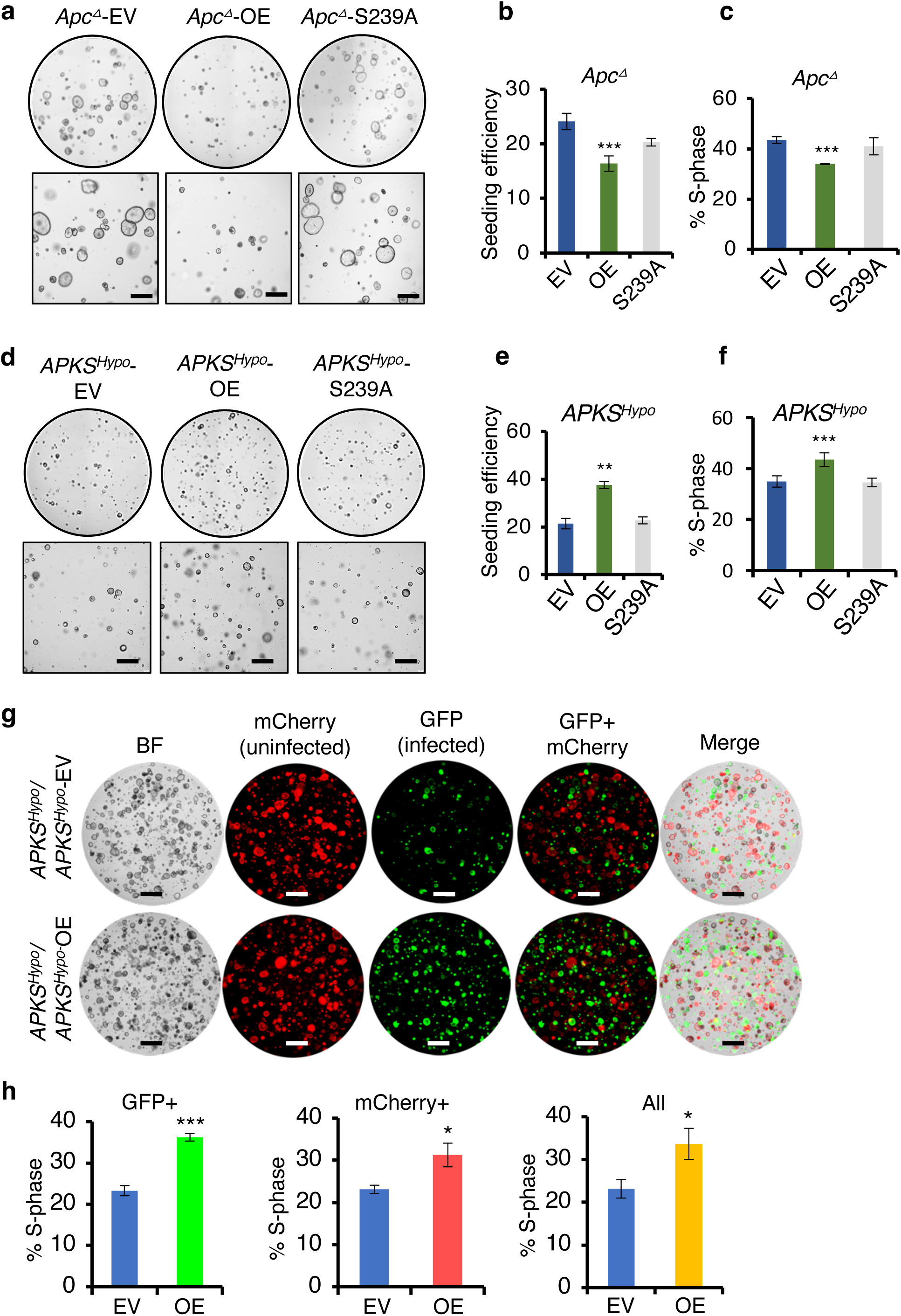
Tumor suppressive and oncogenic functions of NOTUM require catalytic activity and occur via paracrine/juxtacrine signaling. **(a)** Brightfield images of *Apc* mutant tumoroids infected with empty vector (EV), vector expressing wildtype NOTUM (OE), or catalytically dead NOTUM with Ser239Ala mutation (S239A) (n=3). Scale bar: 100 *μ*m. **(b)** Clonal seed efficiency from single cells in (**a**). **(c)** EdU incorporation assays in tumoroids from (**a**). **(d)** Brightfield images of the *APKS* subclone with hypomorphic NOTUM mutation from Fig. S1N/O (*APKS^Hypo^*) infected with empty vector (EV), vector expressing wildtype NOTUM (OE), or catalytically dead NOTUM with Ser239Ala mutation (S239A) (n=3). Scale bar: 100 *μ*m. **(e)** Clonal seed efficiency from single cells in (**d**). **(f)** EdU incorporation assays in *APKS^Hypo^* organoid from (**d**). **(g)** Images of *APKS^Hypo^* tumoroids, with tumoroids infected with an mCherry-expressing vector and co-cultured with tumoroids infected with a vector expressing GFP only (EV) or expressing GFP and NOTUM (OE) (n=3). Scale bar: 100 *μ*m. **(h)** EdU assays in the cultures in (**g**), quantifying the percentage of cells in S phase in the GFP+ or mCherry+ populations, or the overall population. \**P*<0.05, \*\*\**P*<0.001, Student’s *t*-test.

We next asked whether NOTUM’s tumor suppressive and oncogenic activities in *Apc*^△^ and *APKS* tumoroids, respectively, can occur in a paracrine manner. To this end, we GFP-tagged *Apc*^△^ tumoroids overexpressing NOTUM or an empty vector and co-cultured them with unlabeled *Apc*^△^ tumoroids. Both GFP+ cells overexpressing NOTUM and GFP-negative cells (not overexpressing NOTUM) in these cultures proliferated slower than empty vector controls, confirming that NOTUM’s tumor suppressive activity in the absence of functional APC occurs in a paracrine or juxtacrine manner (**Extended Data Fig. 3c, d**). We performed an analogous experiment in *Notum* hypomorphic *APKS* tumoroids, this time tagging half the culture with mCherry and the other half with a vector expressing GFP and NOTUM or a GFP-only control. As with *Apc*^△^ tumoroids, we found that green, NOTUM overexpressing cells were able to drive the proliferation of mCherry+ cells non-cell-autonomously, as well as the cell-autonomous proliferation of GFP+ cells in *APKS* tumoroids (**Fig. 3g, h**). We therefore conclude that the novel oncogenic activity of NOTUM requires catalytic activity and occurs extracellularly.

### *Apc* and *Trp53* inactivation synergize to confer oncogenic activity upon NOTUM

Having established that NOTUM enzymatic activity is tumor suppressive in wildtype organoids and *Apc*^△^ tumoroids, and is oncogenic in *Apc/Trp53* mutant tumoroids (as well as in *APK* and *APKS* tumoroids), we reasoned that a genetic switch must underly the reversal of NOTUM function, and that P53 inactivation may be responsible. *Notum* expression is strongly upregulated upon *Apc* loss, and remains elevated upon accumulation of oncogenic *Trp53*, *Kras*, and *Smad4* mutations (**Fig. 4a)**. In contrast, inactivation of *Trp53* alone had little effect on *Notum* expression (**Fig. 4a**). We assayed the effects of *Notum* LOF and GOF in *Apc*^△^, *Trp53*^△^, and double mutant (*AP*) tumoroids and found, remarkably, that NOTUM activity remains tumor suppressive in both *Apc*^△^ and *Trp53*^△^ tumoroids, but together these mutations synergize to confer oncogenic activity upon NOTUM (**Fig. 4b-f** and **Extended Data Fig. 4 a-c**).

**Fig. 4.**
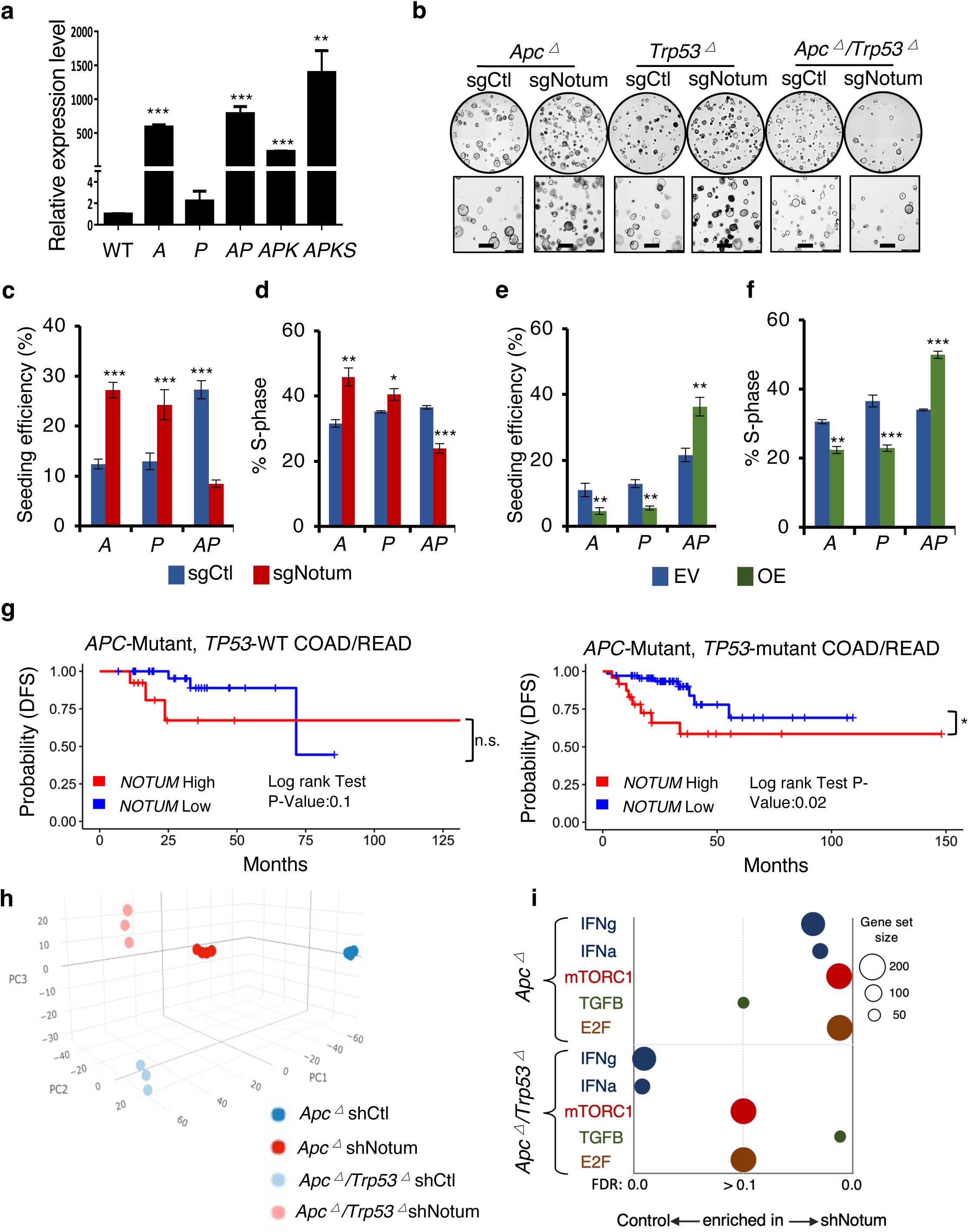
APC and P53 loss synergize to convert NOTUM from a tumor suppressor to an oncoprotein. **(a)** Relative *Notum* transcript levels in WT colon organoids, and tumoroids harboring *Apc* loss of function (*A*), *Trp53* loss of function (*P*), both (*AP*), *AP* with *Kras^G12D^* mutation (*APK*) and *APK* with *Smad4* loss of function (*APKS*) (n=3). **(b)** Brightfield image of *A*, *P* and *AP* tumoroids infected with control sgRNA (sgCtl) or *Notum* sgRNA (sgNotum) (n=3). Scale bar: 100 *μ*m. **(c)** Clonal seeding efficiency of *A*, *P* and *AP* cells from (**b**). **(d)** EdU incorporation assays in *A*, *P*, and *AP* tumoroids from (**b**) (n=3). **(e)** Clonal seeding efficiency of *A*, *P* and *AP* cells infected with NOTUM overexpressing lentivirus (OE) or empty vector control (EV) (n=3). **(f)** EdU incorporation assays in *A*, *P*, and *AP* tumoroids as in (**e**). **(g)** Analysis of disease-free survival (DFS) in colon and rectal adenocarcinoma patients (COAD, READ) for which gene expression data is available in the Cancer Genome Atlas (TCGA). Left panel shows DFS for *APC* mutant, *TP53* wildtype patients (n=46), binned on highest and lowest quartile of *NOTUM* expression (Q1 vs Q4). Right panel shows DFS for *APC/TP53* double mutant COAD/READ patients (n=93), binned on highest and lowest quartile of *NOTUM* expression (Q1 vs Q4). **(h)** Principal component analysis (PCA) of transcriptome profiles from *Apc* mutant or *Apc/Trp53* mutant cultures with or without shRNA-mediated knockdown of *Notum* (n=4). **(i)** Gene set enrichment analysis of transcriptome profiles from (**h**). For all panels: \**P*<0.05, \*\**P*<0.01, \*\*\**P*<0.001, Student’s *t*-test.

Given the striking nature of these findings, we asked whether evidence exists in human colorectal cancers for mutational background-dependent oncogenic NOTUM activity. We pooled patient data from colon and rectal adenocarcinoma (COAD and READ) from the Cancer Genome Atlas (TCGA), then stratified patient disease-free survival based on *NOTUM* expression (top vs. bottom 50% or quartile 1 vs quartile 4) in patients that have *APC* mutations but wildtype *TP53*, or those harboring both *APC* and *TP53* mutations. Consistent with our findings in the mouse tumoroid model, high *NOTUM* expression predicts significant decreases in disease-free survival only in *TP53-*mutant tumors and not in *TP53* wildtype tumors harboring *APC* mutations (**Fig. 4g and Extended Data Fig. 4d**). These differences are likely under-appreciated since the function of wildtype P53 can be impeded in a manner that does not involve mutation (e.g., MDM2 overexpression) ^[20]^. Given that *APC* and *TP53* are the two most frequently mutated genes in colorectal cancer and often coincident in advanced tumors ^[3]^, these findings have far-reaching clinical implications for advanced colorectal adenocarcinoma.

In an effort to delineate the molecular mechanisms underlying *Notum*’s tumor suppressive-to-oncogenic switch downstream *Apc* and *Trp53* mutation, we performed transcriptome profiling of *Apc*^△^ and *AP* cultures, with or without NOTUM LOF (**Fig. 4h**). The transcriptional pathways most differentially affected by NOTUM LOF between *Apc*^△^ and AP double mutant cultures included mTORC1 and E2F signaling (induced by NOTUM LOF in *Apc*^△^ but not *AP*), Interferon a/y signaling (induced by NOTUM LOF in *Apc*^△^, suppressed in *AP*), and TGFβ (induced by NOTUM LOF in *AP*, not in *Apc*^△^) (**Fig. 4i, Extended Data Fig. 4e, and Supplemental Table 2**). These transcriptional pathways reside downstream of cell-surface receptors, many of which are known to be influenced by the activity of Glypicans, a family of proteoglycans that can be cleaved from the cell surface by NOTUM activity ^[8, 10]^. For example, Glypicans have been implicated in the regulation of WNT, TGF/3, IGF, FGF, VEGF, Hh, HGF, and BMP receptor activity, primarily in studies carried out in *Drosophila* ^[8, 21–32]^. In mammals, how each of the six paralogous orthologs of *Drosophila* Glypicans (mammalian Gpc1-6) influence activity of these receptors and in which context remains poorly understood.

We examined the expression of the mammalian Glypicans in mouse tumors, mouse tumoroids, human tumors, and human tumoroids and found *Gpc/GPC1* and *Gpc/GPC4* consistently to be the dominantly expressed family members in CRC (**Extended Data Fig. 5a-d**). Given the models established in the literature, we hypothesized that upon cleavage of their glycosyl-phosphatidylinositol (GPI) anchor by NOTUM, GPCs released from the cell surface may either be neutralized or negatively regulate receptor activity due to their ability to compete with cell surface-bound receptors for their cognate ligands.

### The tumor suppressive versus oncogenic functions of NOTUM can be accounted for by its cleavage of cell surface glypicans

We initially tested whether genetic inactivation of *Gpc1/4* could phenocopy NOTUM activity. Interestingly, while GPC1/4 protein levels were both readily detectible in both *Apc*^△^ and *AP* tumoroids (**Extended Data Fig. 5e**), GPC1 exhibited oncogenic activity only in *Apc*^△^ cultures, with no effect in *AP* tumoroids.

Conversely, GPC4 exhibited tumor suppressive activity only in *AP* tumoroids, with no effect in *Apc*^△^ tumoroids (**Fig. 5a-b, Extended Data Fig. 5f-h**). *Gpc6*, whose expression was low but detectable in this system, had little to no effect in either genotype (**Fig. 5a-b**). Next, to link NOTUM function, GPC protein, and signaling pathway activity, we knocked down *Notum* in *Apc*^△^ and AP tumoroids and examined pathway activity and cell-associated GPC protein levels. *Notum* knockdown in *Apc*^△^ tumoroids lead to increased mTORC1 signaling coupled with an accumulation of GPC1 protein, with no apparent changes to TGFβ pathway activity (**Fig. 5c**). In contrast, *Notum* knockdown in *AP* tumoroids lead to increased TGFβ pathway activity and an accumulation of GPC4 protein, with no apparent effect on mTORC1 activity (**Fig. 5d**). These experiments assayed GPC protein levels in whole cell lysates and support a model whereby *Notum* loss of function prevents cleavage of GPCs from the cell surface leading to their accumulation, as observed in Drosophila studies ^[8, 10]^. To further probe this possibility, we examined levels of free N-terminal GPC1 and GPC4 in the media supernatant of *Apc*^△^ and *AP* tumoroids after NOTUM inhibition. NOTUM inhibition in *Apc*^△^ tumoroids increased cell-associated GPC1 relative to GPC1 in the supernatant (**Fig. 5e**). Conversely, *Notum* loss of function in AP tumoroids led to increased cell-associated GPC4 concomitant to decreased GPC4 in the supernatant (**Fig. 5e**). Interestingly, NOTUM protein itself was associated with the cell lysate and not supernatant, a finding that, in conjunction with the extracellular enzymatic effects of NOTUM observed in **Fig. 3g** and **Extended Data Fig. 3**, suggests that NOTUM is physically associated with the extracellular cell surface (**Fig. 5e**).

**Fig. 5.**
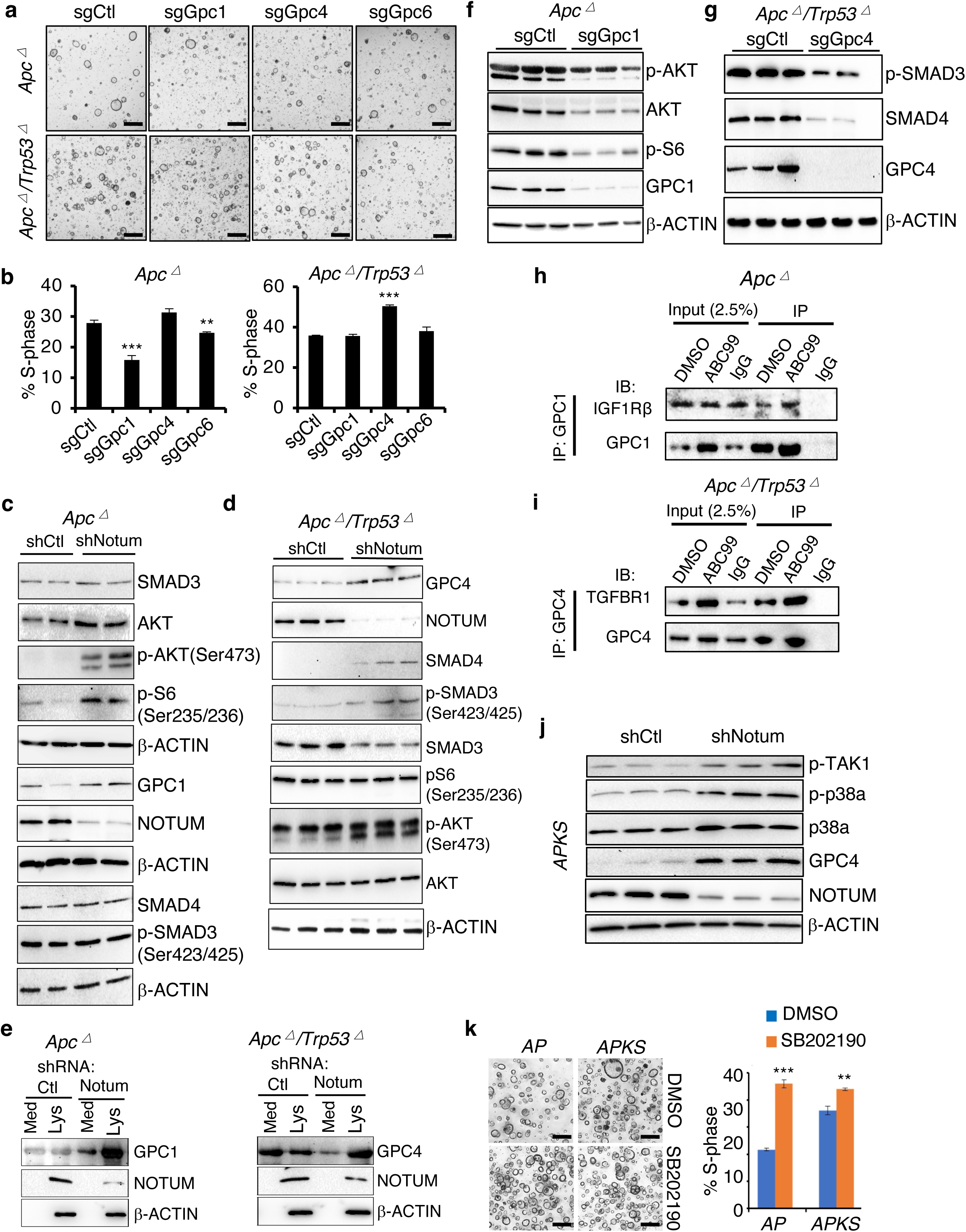
Glypicans mediate the differential effects of NOTUM activity in *Apc* mutant versus *Apc/Trp53* mutant tumoroids. **(a)** Brightfield image of *Apc* mutant tumoroids (*A*) and *Apc/Trp53* double mutant tumoroids (*AP*) infected with control sgRNA (sgCtl), *Gpc1* sgRNA (sgGpc1), *Gpc4* sgRNA (sgGpc4), *Gpc6* sgRNA (sgGpc6) (n=3). Scale bar: 100 *μ*m. **(b)** EdU assays quantifying fraction of cells in S-phase from cultures shown in (**a**) (n=3). **(c)** Western blotting in *Apc* mutants infected with control shRNA (shCtl) or *Notum* shRNA (shNotum). β-ACTIN was used as a loading control. Note multiple β-ACTIN blots, one for each of three identical replicate blots. **(d)** Western blotting in *Apc/Trp53* mutant tumoroids infected with control shRNA (shCtl) or *Notum* shRNA (shNotum). β-ACTIN was used as a loading control. **(e)** Western blotting for GPC1 (left) and GPC4 (right) using N-terminal antibodies recognizing the extracellular domain of indicated glypicans in the media supernatant (Med) or cell lysate (Lys) of *Apc* mutant or *Apc/Trp53* double mutant tumoroids infected with control shRNA (shCtl) or *Notum* shRNA (shNotum). β-ACTIN was used as a loading control. **(f)** Western blotting in *Apc* mutant tumoroids infected with control sgRNA (sgCtl) or *Gpc1* sgRNA (sgGpc1). β-ACTIN was used as a loading control. **(g)** Western blotting in *Apc/Trp53* mutant tumoroids infected with control sgRNA (sgCtl) or *Gpc4* sgRNA (sgGpc4). β-ACTIN was used as a loading control. **(h)** Co-immunoprecipitation of IGF1Rb with anti-GPC1 antibody in *Apc* mutant tumoroids in the presence or absence of the small molecule NOTUM inhibitor ABC99. **(i)** Co-immunoprecipitation of TGFβR1 with anti-GPC4 antibody in *Apc/p53* double mutant tumoroids in the presence or absence of the small molecule NOTUM inhibitor ABC99. **(j)** Western blotting assessing non-canonical TGFβ-TAK1-p38α pathway activity in *APKS* tumoroids in response to NOTUM inhibition. β-ACTIN was used as a loading control. **(k)** *AP* and *APKS* tumoroids treated with the p38 inhibitor SB202190, with EdU-based quantification of S-phase (n=3). Scale bar: 100 *μ*m. For all panels: \*\**P*<0.01, \*\*\**P*<0.001, Student’s *t*-test.

To further test whether the effects of GPC manipulation are consistent with a model whereby NOTUM cleaves GPC1 and 4 from the surface of *Apc*^△^ tumoroids and AP tumoroids, respectively, we analyzed the effects of GPC inactivation on the molecular pathways differentially effected by NOTUM activity. Indeed, GPC1 inactivation in *Apc*^△^ tumoroids inhibited mTORC1 activity, and GPC4 inactivation in *AP* tumoroids inhibited TGFβ activity, consistent with the effects of NOTUM manipulation (**Fig. 5f, g**) and supporting a model where NOTUM cleavage of GPCs from the cell surface results in their loss of function in potentiating these signaling pathways. Finally, we sought to confirm direct interaction between GPC1 or GPC4 and the receptors upstream of the affected signal transduction pathways. Immunoprecipitation of GPC1 in *Apc*^△^ tumoroids co-immunoprecipitated IGF1Rβ, an IGF receptor upstream of mTORC1 activation implicated in colon cancer and chemotherapeutic resistance ^[33, 34]^ (**Fig. 5h**). In *AP* tumoroids, immunoprecipitation of GPC4 co-immunoprecipitated TGFβR1 upstream of the TGFβ pathway (**Fig. 5i**). Addition of ABC99, a small molecule inhibitor of NOTUM catalytic activity ^[9, 38]^, appeared to potentiate these interactions (**Fig. 5h, i**).

Taken together, these findings support a model whereby NOTUM exerts its tumor suppressive effects in *Apc*^△^ cells by suppression of oncogenic GPC1 activity upon IGF1Rβ upstream of mTORC1 activation. In contrast, upon *Trp53* inactivation NOTUM exerts oncogenic effects via cleavage of tumor suppressive GPC4 from the cell surface to inhibit TGFβR1 - TGFβ activity. However, we also observed clear oncogenic effects of NOTUM in the highly aggressive *APKS* tumoroids and tumors, where the canonical TGFβ pathway is not functional due to inactivating mutations in *Smad4* (an established suppressor of colon cancer metastasis) ^[35, 36]^. We therefore interrogated the effects of NOTUM modulation on non-canonical TGFβ pathway activity mediated through TAK1-p38α signal transduction. Consistent with our findings in *AP* tumoroids, NOTUM inhibition resulted in increased GPC4 in APKS cell lysate (**Fig. 5j**). Further consistent with NOTUM’s effects as a negative regulator of TGFβ signaling, *Notum* inhibition in *APKS* tumoroids resulted in increased phosphorylation of TGFβ-Activated Kinase (TAK1) and downstream p38α phosphorylation (**Fig. 5j**). The role of p38α activity in cancer is pleiotropic and context-dependent, including evidence for both oncogenic and tumor suppressive effects (the latter through cell cycle arrest, differentiation, senescence, and activation of apoptotic pathways, particularly in KRAS-activated cancers) ^[37–39]^. We observe that p38α inhibition in both *AP* and *APKS* tumoroids increased proliferation and inhibited cell death (**Fig. 5k and Extended Data Fig. 5i**). In *APKS* tumoroids, NOTUM inhibition induced apoptosis, and this was reversed by concomitant inhibition of p38α (**Extended Data Fig. 5i**). These findings support the notion that NOTUM may exert oncogenic effects, at least in part, via inhibition of this tumor suppressive activity of the non-canonical TGFβ pathway.

### Pharmacological NOTUM inhibition is efficacious in a pre-clinical animal model of metastatic colon cancer

Ultimately, we asked whether small molecule inhibition of NOTUM represents a viable therapeutic strategy in advanced colorectal cancers harboring inactivating mutations in *Apc* and *Trp53*. ABC99 is a potent and selective small molecule inhibitor of NOTUM with *in vivo* efficacy ^[19, 40]^. We initially confirmed that ABC99 treatment phenocopies genetic inhibition of NOTUM in *Apc*^△^ tumoroids and highly metastatic *APKS* tumoroids (**Fig. 6a, b and Extended Data Fig. 6a, b**). Next, we orthotopically implanted *APKS* tumoroids into the colonic mucosa of syngeneic mice via endoscope-guided injection and allowed tumors to engraft and grow for one month, after which mice were randomly assigned to a vehicle control or ABC99 group. An absence of statistical difference in tumor size was confirmed between the two randomized groups at baseline using bioluminescence imaging (BLI) and endoscopy (**Extended Data Fig. 6c, d**). Mice were then treated with ABC99 or vehicle control for an additional four weeks. ABC99 treatment largely arrested primary tumor growth, resulting dramatic decreases in both BLI and tumor weight relative to control animals (**Fig. 6c-e**). Remarkably, NOTUM inhibition with ABC99 nearly completely inhibited the formation of liver and lymph node metastases in this highly aggressive model of colorectal adenocarcinoma (**Fig. 6f, g and Extended Data Fig. 6e, f**). Relative to vehicle-treated controls, ABC99-treated primary and rare metastatic lesions exhibited notable inhibition of cell proliferation at the termination of the experiment (**Fig. 6h**). Given these striking findings, we generated another cohort of mice with orthoptic *APKS* tumors, allowed them to engraft for one month as before, then began ABC99 or vehicle control treatment and monitored survival for two months. Remarkably, while the majority of vehicle control-treated mice became moribund over this 3-month time course, none of the ABC99-treated cohort succumbed to disease for the duration of this experiment (**Fig. 6i**).

**Figure 6.**
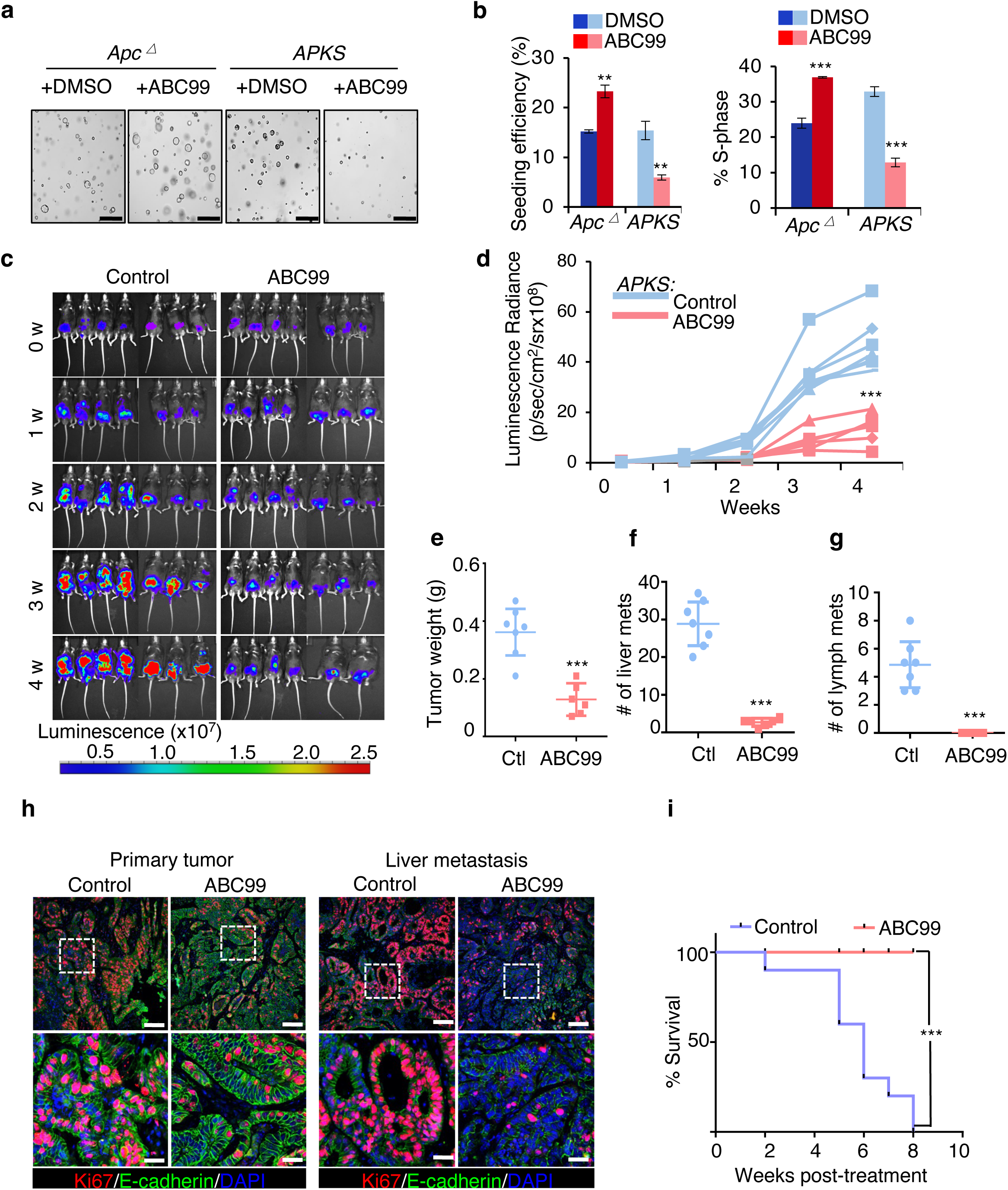
Small molecule inhibition of NOTUM inhibits tumor growth and metastasis in a mouse model of colorectal cancer. **(a)** Brightfield image of *Apc* mutant and *APKS* tumoroids treated with the small molecule inhibitor of NOTUM, ABC99, or vehicle control (DMSO) (n=3). Scale bar: 100 *μ*m. **(b)** Single cell clonal seeding efficiency and EdU assay quantifying fraction of cells in S phase from tumoroids in (**a**) (n=3). **(c)** Bioluminescent imaging (BLI) of mice after endoscope-guided orthotopic implantation of *APKS* tumoroids into the colonic mucosa. Mice were treated with ABC99 or vehicle (corn oil) control after 4 weeks of engraftment and tumor growth. Mice are treated for indicated number of weeks (week 0 = 4 weeks post-implantation) (n=7 mice). **(d)** Quantification of luminescence radiance from (**c**) at indicated timepoints (n=7 mice). **(e-g)** Quantification of primary tumor weight (**e**), number of macrometastases in the liver (**f**), and number of lymph node metastases (**g**) in mice from (**c**) at the termination of the experiment (8 weeks total, 4 weeks post-treatment). **(h)** Immunofluorescence staining for KI67 and E-CADHERIN in primary tumor and liver metastases from tumors in (**c**) (n=7 mice). Nuclei are visualized by DAPI staining. Scale bar of top panels: 200 *μ*m. Lower panels: 50 *μ*m. **(i)** Kaplan-Meier survival analysis for mice harboring orthotopic *APKS* tumors as in (**c**), treated with ABC99 or vehicle (corn oil) control for 8 weeks following an initial 4 weeks period of engraftment and tumor growth (total experimental time course of 3 months) (n=10 mice). For all panels, \*\**P*<0.01, \*\*\**P*<0.001, Student’s *t*-test.

Finally, we asked whether NOTUM inhibition with ABC99 resulted in similar efficacy in human CRC tumoroids. Treatment of *APC/TP53* double mutant human colon tumoroids with ABC99 inhibited clonal tumoroid initiation capacity and proliferation, indicating that small molecule inhibition of NOTUM is also a viable strategy in human cells (**Extended Data Fig. 6g-i**). Ultimately, these findings identify NOTUM as a potentially tractable therapeutic target in advanced colorectal adenocarcinoma harboring inactivating mutations in both APC and TP53.

## Discussion

Advanced colorectal cancer remains a leading cause of cancer-related deaths globally, with few efficacious therapeutic options available once the primary tumor has disseminated. While it is long- and well-established that inactivation of the APC tumor suppressor is likely the initiating event in the vast majority of CRC, how the initial events downstream of *Apc* inactivation influence later stages of tumorigenesis after accumulation of additional oncogenic mutations remains unclear. Recent data, however, suggest that APC inactivation is not only required for tumor initiation, but also for tumor maintenance in more advanced disease. Reactivation of APC in *Apc/Kras/Trp53* triple mutant mouse tumors is sufficient to cause tumor regression and restore seemingly normal crypt-villus homeostasis despite the continued presence of oncogenic KRAS and absence of P53 function ^[14]^. Activation of the extracellular palmitoleoyl-protein carboxylesterase NOTUM has long been observed downstream of APC inactivation, and *Notum* is recognized as a direct β-Catenin transcriptional target gene ^[16]^.

In mammals, NOTUM has recently gained attention as a negative regulator of canonical WNT pathway activity. While nearly undetectable in the epithelium of young mice, NOTUM expression increases as a function of age, where it has been linked to decreases in WNT and intestinal stem cell activity ^[19]^. Remarkably, in aged animals, systemic, small molecule inhibition of NOTUM activity restores stem cell function, presumably through potentiation of canonical WNT activity ^[19]^. In mammals, this negative regulation of WNT activity by NOTUM is attributed to its ability to cleave the palmitoleate tail from WNT ligands, thus rendering them unable to interact with Frizzled (FZD) receptors in order to transduce the WNT signal ^[11]^.

In the context of intestinal tumorigenesis, the rapid and robust induction of *Notum* expression downstream of APC inactivation might be viewed as a futile attempt of the transformed cell to attenuate the oncogenic consequences of constitutive β-Catenin transcriptional activity. Indeed, a recent study put forth the hypothesis that the robust induction of NOTUM in an intestinal epithelial stem cell that has inactivated APC gives that *Apc^−/−^* clone a competitive advantage over its *Apc*-wildtype neighbors via NOTUM-mediated juxtacrine inhibition of canonical WNT signaling ^[12, 13]^. On the surface, these findings appear contradictory with our current observations. However, these findings can potentially be reconciled if the anti-proliferative effects of NOTUM are stronger on wildtype versus *Apc^−/−^* stem cells, which is reasonable to assume given the exquisite dependence of wildtype stem cells on WNT ligand/receptor interactions. In this context, a *Notum*-null clone would have a proliferative advantage over its wildtype neighbors. However, upon reaching some critical size at which the adenomatous lesion is no longer constrained by neighboring wildtype epithelial cells (as in our orthotopic implantation model), the cell-autonomous tumor suppressive role of *Notum* in an *Apc^−/−^* clone, which has previously not been addressed, becomes evident. The unexpected finding that NOTUM retains potent tumor suppressive activity in adenomatous *Apc^−/−^*cells is not readily explicable by the extracellular mechanism(s) through which NOTUM antagonizes WNT pathway activity. Similarly, our observation that *Notum* becomes an obligate oncogene once these tumors progress to adenocarcinomas with subsequent P53 inactivation is also not explicable by its function as a WNT antagonist.

Instead, a body of literature primarily derived from studies in *Drosophila melanogaster* point to a potentially much broader role for NOTUM in the regulation of numerous signal transduction cascades via its ability to cleave glypicans from the cell surface. Glypicans are a family of proteoglycans, comprising two members in drosophila (*Dally* and Dally-like-protein, *Dlp*), that have expanded to six paralogous orthologs in mammals (*Gpc1-6*) ^[41]^. Dally and Dlp have been implicated in the regulation of most known signal transduction pathways, including WNT, TGFβ, IGF, FGF, VEGF, Hh, HGF, and BMP ^[8, 21–32]^. Most current models for Glypican function indicate that these proteoglycans, when anchored to the cell surface, act to potentiate receptor-ligand interactions. NOTUM can cleave GPCs at their glycosyl-phosphatidylinositol linkage to the cell surface ^[8, 10]^, and our findings suggest that GPC1 and GPC4 are key family members in colorectal cancer. Once cleaved from the cell surface, N-terminal GPCs can now act as competitive inhibitors of the same signaling pathways they potentiate when surface-bound, via ligand competition. However, some evidence also suggests that surface-bound GPCs may negatively regulate receptor-ligand interactions. For example, studies in mammalian cells indicate GPC3 that can compete for Hedgehog ligand binding with the Patched receptor ^[42]^, although this contrasts with evidence in Drosophila that Dlp is an agonist of Hedgehog pathway activity ^[43]^.

Ultimately, the function of mammalian Glypicans is poorly understood, and their role in colorectal cancer remains unstudied. In the current study, gene expression changes downstream of *Notum* loss-of-function appear context-dependent in *Apc*-mutant versus *Apc/Trp53* double mutant cells, with clear induction of mTORC1 activity in response to *Notum* inactivation in *Apc*-mutant cells, and activation of TGFβ activity in *Apc/Trp53* double mutants. This finding is consistent with the observed functions of GPC1 and GPC4 in *Apc*-mutant and *Apc/Trp53* double mutant cultures, respectively, as well as with the model that Glypicans potentiate signal transduction at the receptor-ligand level, and the negative regulation of this interaction by NOTUM (**Fig. 7**). While these downstream pathways likely contribute to the tumor suppressive and oncogenic effects of NOTUM activity in *Apc*-mutant and *Apc/Trp53* double mutant cultures, respectively, there remains a possibility that GPC1 and GPC4 also act upon additional signaling pathways, and that additional Glypicans may be affected by NOTUM activity during the ontogeny of colorectal adenocarcinomas. Nonetheless, despite the potential for pleiotropic functions of NOTUM in colon cancer, the clear and compelling finding that extracellular enzymatic activity of NOTUM is required for adenocarcinoma maintenance and progression, coupled with the finding that small molecule inhibition of NOTUM activity is sufficient to arrest tumor progression and metastasis in immune-competent mouse models of invasive colon adenocarcinoma indicate that NOTUM is a potentially tractable therapeutic target in this deadly disease.

**Fig. 7:**
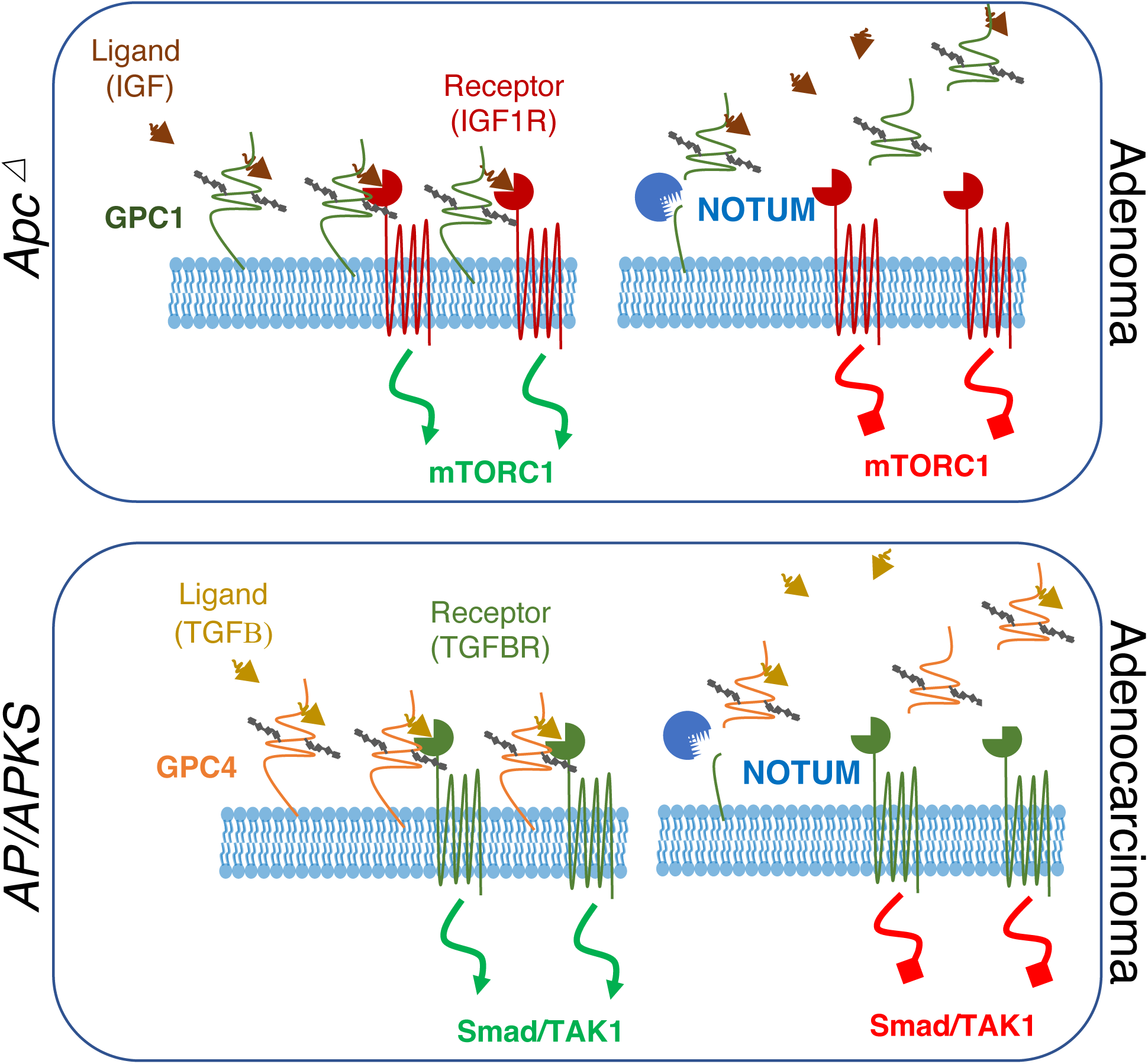
Model for NOTUM tumor suppressive-to-oncogenic switch during the adenoma-adenocarcinoma transition. In early adenomas resulting from Apc inactivation, NOTUM exerts tumor suppressive activity through cleavage of oncogenic Glypican 1 from the cell surface. Here, Gpc1 acts as an oncogene by potentiating IGF1R activity upstream of mTORC1, and Notum antagonizes this oncogenic axis (TOP). In contrast, upon transition to adenocarcinoma with inactivation of p53 (and upon accrual of additional driver mutations), NOTUM functions as an obligate oncogene by inhibiting tumor suppressive TGFβR activity via inactivating cleavage of Glypican 4 from the cell surface, which normally potentiates TGFβR activity upstream of TAK1/Smad signaling.

## Methods

### Mouse models

All procedures involving mice were reviewed and approved by the Institutional Animal Care and Use Committee and were in accordance with the guidelines set forth in the Guide for the Care and Use of Laboratory Animals of the National Research Council of the National Institutes of Health. All animals were maintained on a C57BL/6 genetic background. KrasLSL-G12D (JAX strain 008179), C57BL/6J (JAX strain 000664) mice were obtained from The Jackson Laboratory. After orthotopic tumoroid injection, tumors were allowed to engraft for 4 weeks, after which mice were injected intraperitoneally daily with ABC99 at (Sigma-Aldrich # SML2410) suspended in corn oil (Sigma-Aldrich # C8267) at a concentration of 5 mg/ml and 50 ul per 25 g of body weight.

### Organoid culture

#### Mouse organoid culture

Male C57BL/6J or KrasLSL-G12D mice were euthanized by carbon dioxide followed by cervical dislocation at around the age of 48 days. Colon crypts were isolated and cultured as described previously ^[44]^. Organoids were seeded in plate mixed with growth factor reduced Matrigel (Corning #356231). Culture media was made of Advanced DMEM/F-12 (Thermo Fisher Scientific #12634010) containing B27(Gibco # 17504-044), N2 (Invitrogen # 17502-048), 2mM GlutaMAX (Thermo Fisher Scientific # 35050061), 10mM HEPES (Thermo Fisher Scientific # 15630-080), 1mM N-acetylcysteine (Sigma-Aldrich# A9165), supplemented with 50 ng/ml mEGF (Invitrogen # PMG8041) and 1x conditioned media (derived from L-WRN cells).

#### Human patient-derived organoid culture

0.5–2 cm^3^ portions of fresh CRC surgical specimens were obtained from patients undergoing surgical resection for treatment-naïve colorectal cancer. All samples were obtained from patients who provided informed consent before surgery. Cancer and normal epithelia tissue were isolated and cultured according to previously reported protocols ^[45]^. Briefly, dissected the mucosal layer off from the whole normal tissue, and then digested with 5 mM EDTA 4 °C for 20 min on the roller. Released the crypts by squeezing the mucosal layer hard and collected the released crypts back by spin down at 300g for 5min. Then seeded with Matrigel into 24-well plates. For cancer tissue, digested with collagenase Type IV (Gibco# 17104019, 200 units in 5 ml Dispase) 37 °C for 30 min. After digesting into clumps of cells, the sample was filtered through 70 µm cell strainer and seeded with Matrigel into 24-well plates. The culture of human organoid medium was as follows: Advanced DMEM/F12, 10 mM HEPES, 1× Penicillin-Streptomycin (Invitrogen # 15140122), 2 mM GlutaMAX, 1× N2 supplement, 1× B27 supplement, 1 mM N-acetyl-cysteine, 10 nM human gastrin I (Sigma-Aldrich # G9145), 50 ng/ml hEGF (R&D Systems#236-EG-200), 100 ng/ml Noggin(R&D Systems # 6057-NG-100), 500 nM A83-01 (Tocris # 2939), 10uM SB202190 (Sigma-Aldrich # S7067), 30ug/ml humanWnt-3a (ProSpec # CYT-861), 1mg/ml R-spondin (R&D Systems #4645-RS-025), and 10 uM Y-27632(Selleck Chemicals # S1049).

#### Human iPS-derived colon organoid culture

Wildtype human iPSCs. were maintained on plates coated in growth factor reduced Matrigel (Corning) and cultured in StemMACS™ iPS-Brew XF medium (Miltenyi Biotec) supplemented with 1% Pen-Strep (Gibco). Cells were cultured in a 5% CO_2_, 5% O_2_, and 37°C incubator and passaged every 4-5 days in small clusters using StemMACS Dissociation reagent (Miltenyi Biotec). Colon organoids were differentiated as previously described^[46]^ with minor modifications. Briefly, cells were washed with PBS, dissociated into single cells using warmed Accutase (STEMCELL Technologies), and seeded at a density of 500,000 cells per well of a matrigel-coated 6-well plate in iPS-Brew XF supplemented with 2 uM Thiazovivin ROCK inhibitor (Cayman Chemical). Definitive endoderm was derived using STEMdiff™ Definitive Endoderm Kit (STEMCELL Technologies) according to the manufacturer’s instructions. Following definitive endoderm differentiation, cells were subjected to hindgut differentiation by treatment with 500 ng/ml FGF4 (Peprotech) and 3 μM CHIR99021 in RPMI supplemented with 1X B27 supplement (Gibco), 1X GlutaMAX (Gibco), and 1% Pen-Strep (Gibco). Hindgut medium was refreshed daily for a total of 4 days. From day 8 onward, cells were cultured in colonic differentiation medium containing Advanced DMEM/F12 (Invitrogen) supplemented with 1X B27 supplement (Gibco), 1X GlutaMAX (Gibco), 1% Pen-Strep (Gibco), 3 μM CHIR99021 (Cayman Chemical), 300 nM LDN193189 (Cayman Chemical) and 100 ng/ml EGF (Peprotech). 2D cultures were monitored daily for the next 10-12 days for the emergence of 3D spheroid structures. Where necessary, 3D spheroids were aided in detaching from the monolayer by gentle pipetting. Spheroids were collected and embedded in growth factor reduced Matrigel (Corning) and overlaid with colonic differentiation medium. Organoids were passaged every 10-12 days by mechanical dissociation. Mature organoid cultures were validated by confirming the presence of terminally differentiated colonic cell types by IHC and quantifying the expression of colonic marker genes by qRT-PCR compared to undifferentiated iPSC controls. To generate APC and P53 knockout (*AP*) CRISPR tumoroids, gRNA sequences targeting *Apc* (5-CAACTTCTGGTAATGGTC-3) or *Trp53* (5-AATGAGGCCTTGGAACTCA-3) were cloned into the Px330 vector (Addgene plasmid #42230). Organoids were removed from Matrigel using Dispase II (Gibco), dissociated into single cells using warmed TrypLE Express (Gibco), and resuspended in colonic differentiation medium without antibiotic. CRISPR vectors were transfected into cells using Lipofectamine Stem (Invitrogen) according to the manufacturer’s instructions. Organoid transfection plates were then centrifuged at 600 g for 1 hr at 32°C and incubated in a 37°C incubator for an additional 3-4 hrs to allow transfection. Organoid cells were collected, centrifuged, replated in serial dilutions of growth factor reduced Matrigel, overlaid with colonic differentiation medium supplemented with 2 uM Thiazovivin (Cayman Chemical) and allowed to recover for 48 hours. Two days after transfection, organoid medium was replaced with AP selection medium (colonic differentiation medium lacking CHIR99021 and supplemented with 5 uM Nutlin-3 (Cayman Chemical) to select for organoids with loss of *APC* and *TP53* function, respectively). Selection medium was refreshed every other day for 10-12 days. After 12 days, individual *APC* and *TP53* inactivating mutations were confirmed by Sanger sequencing.

### Endoscope-guided orthotopic transplantation

Approximately 8-week-old male C57BL/6J mice from the Jackson laboratory were used for orthotopic transplantation experiments. Luciferase-expressing organoids were isolated from Matrigel using Cell Recovery Solution (Corning # 354253) incubated on ice for 30 min. Cell clusters (equivalent to 2 × 10^4^cells for each mouse) were resuspended in cold D-PBS with 0.5% BSA and 10% Matrigel and then transplanted into the colon mucosa of recipient mice by optical colonoscopy using a custom injection needle (Hamilton Inc., 33 gauge, small Hub RN NDL, 12 inches long, point 4, 45-degree bevel, catalog #7803-05), a 100µl syringe (Hamilton Inc., part number 7656-01), a transfer needle (Hamilton Inc., part number 7770-02), and a colonoscope with integrated working channel (Richard Wolf 1.9 mm/9.5 French pediatric urethroscope, part number 8626.431). 70 µl of cell suspension was delivered to the colon mucosa. One injection was performed per mouse. Mice underwent colonoscopy 2-8 weeks following organoid transplantation to assess tumor development. We quantified tumor size in situ by photographing them and measuring the percent of the lumen occluded by tumor. We measured the tumor size relative to the luminal diameter and recorded as the percent tumor occlusion of the lumen. The change in percent tumor occlusion relative to the baseline measurement was calculated every two weeks and averaged within the two groups. At the experimental endpoint, tumor weights were recorded. We also used an IVIS imaging system to monitor tumor development in vivo once a week during treatment with ABC99 or vehicle control until the experimental endpoint 8 weeks post-implantation.

### Generation of vectors and sgRNA cloning

The pX330-U6-Chimeric_BB-CBh-hSpCas9 (Addgene#42230), LentiCRISPR v2 expression vector (Addgene# 52961), LentiCas9-Blast (Addgene# 52962) and LRG (Addgene# 65656) which expressed sgRNA and/or Cas9 nuclease were used to generate gene-specific sgRNA vectors. The pLKO_TRC005 (RNAi Consortium) was used to deliver shRNA. PUltra (Addgene# 24129) was used to cDNA expression. For sgRNA cloning, pX330 was digested with BbsI (NEB # R3539S) and ligated with a BbsI-compatible annealed sgRNA oligos targeting exon 16 in *Apc* (target sequence GTAATGCATGTGGAACTTTG<u>TGG</u>, PAM sequence underlined), or exon 4 in *Trp53* (target sequence <u>CCC</u>TCCGAGTGT CAGGAGCTCC, PAM sequence underlined), or exon 2 in *Smad4* (target sequence GATGTG TCATAGACAAGGT<u>GGG</u>, PAM sequence underlined). Lenti-CRISPR v2 and LRG were digested with BsmBI (NEB#R0739S) and ligated with a *BsmBI*-compatible annealed sgRNA oligos targeting exon 2 in *Notum* (target sequence AGGAAGACCTGGAGGCGCCG<u>TGG</u>, PAM sequence underlined), or exon 1 in *Gpc1* (target sequence CGCTGGTCGTCTGCGCCCGC<u>GGG</u>, PAM sequence underlined), or exon 1 in *Gpc4* (target sequence CGACGTCTCTACGTGTCCAA<u>AGG</u>, PAM sequence underlined), or exon 2 in *Gpc6* (target sequence CCGAGCCTTCACGTCCGCCG<u>AGG</u>, PAM sequence underlined). pLKO_TRC005 was digested with NdeI (NEB# R0111S) and EcoRI (NEB#R3101) and ligated with shNotum oligo targeting exon 10 (target sequence GCCAGTGGCTATACATCCAGA).

For Notum overexpression, Notum-F ATATCTAGAATGGGAGGAGAGGTGCGCGTGCT and Notum-R CGCGGATCCCTAGTTCCCATTACTCAGCATCCCTA were used to clone mouse Notum from colon organoid cDNA. The PCR product and pUltra were digested with XbaI (NEB# R0145S) and BamHI (NEB# R3136T) and then ligated and sequence validated.

### Murine intestinal organoid culture and genome engineering

Murine colon organoids were cultured as previously reported^[44]^ and briefly summarized here. The colons of Kras-LSL-G12D-LSL (The Jackson Laboratory, stock no: 008179) or C57BL/6J (The Jackson Laboratory, stock no: 000664) mice were removed, washed with cold D-PBS multiple times until clean, and digested with 5 mM EDTA (Invitrogen # 15575-020) for 20 min in cold room. Crypts were released under a dissection microscope by squeezing the mucosal layer and then filtered through 70-µm cell strainer to remove tissue fragments. Crypts were counted and spun down at 300 g for 5 min. Crypts were embedded in Matrigel and seeded into 24-well plates (Olympus # 25-107) at a density of ∼500 crypts in 50 µl total volume per well, and incubated with basal medium (Advanced DMEM/F12, 10 mM HEPES, 1× Penicillin-Streptomycin, 2 mM GlutaMAX, 1× N2 supplement, 1× B27 supplement, 1 mM N-acetyl-cysteine), supplemented with 50 ng/ml mEGF (Invitrogen # PMG8041) and 1x conditioned media (derived from L-WRN cells). Cultures were treated with 10 ul Y-27632 (Rho Kinase inhibitor, Selleck Chemicals # S1049) for 2 days after seeding to prevent cell death by anoikis.

CRISPR/Cas9 editing of organoids was performed in two-dimensional culture. 24-well plates were coated with diluted Matrigel (Matrigel: D-PBS=1:49) and incubated for 2 hours at 37°C. Murine colon organoids were gently dissociated into single cells using 1× TrypLE (Thermo Fisher Scientific # 12605010) at 37 °C for 5 min and then seeded onto the 2D plate. *APKS* organoids were made as follows. The KrasG12D mutation was activated by Cre-mediated deletion of the stop cassette at the Kras-LSL-G12D targeted allele by transfection of a plasmid expressing Cre recombinase along with a pPGK-Puro plasmid (Addgene#11349) using Lip2000 (Thermo Fisher Scientific #11668019) followed with 10 ug/ml puromycin selection for 3 days. The *Apc*, *Trp53* and *Smad4* mutations were introduced by CRISPR-Cas9 editing. Specifically, sgRNAs of *Apc*, *Trp53* and *Smad4* were cloned into PX330 plasmid then transiently transfected into organoids. One week after transfection, the organoids with *Apc*, *Trp53* and *Smad4* mutations were selected by removing L-WRN conditioned media and adding 10 µm Nutlin-3 (an Mdm2 inhibitor enabling functional selection for *Trp53* loss of function) (Cayman Chemical Company #10004372-1). 10 subclones were picked from the engineered bulk organoids, conditional PCR and Sanger sequencing were used to verify the mutations in each subclone. The primers that used to do PCR and Sanger sequencing are listed in **Supplementary table 3**.

Individual *Apc* mutant (*A*), *Trp53* mutant (*P*), and *Apc/Trp53* double mutant (*AP*), organoids were generated by seeding normal colon organoids into 2D and transiently transfected with sgRNAs of *Apc* or *Trp53* or both, as described above. 2 days after transfection, organoids with *Apc* loss of function were selected by removing L-WRN conditioned media and adding 100 ng/ml Noggin, while organoids with Trp53 mutation were selected by adding Nutlin-3 (10 um/ml). Subclones were isolated from the engineered bulk culture and targeted loci PCR amplified and Sanger sequenced to verify the mutations in each subclone.

### Lentiviral infection

The viral vectors were Maxi-prepped using ZymoPURE II™ Plasmid Maxiprep (Zymo Research#D4202) and then transfected into HEK293T cells with packaging plasmids (psPAX2 and pMD2G, Invitrogen). The viral supernatant was collected after 48 hours and 72 hours and passed through a 0.45 µm filter. Lenti-X™ Concentrator (Clontech # 631231) was added to the viral supernatant to concentrate virus. Organoids were dissociated into single cells and seeded into 2D plates and infected with lentivirus plus 1 ug/ml polybrene (Sigma-Aldrich# TR-1003-G). Three days after the infection, organoids were selected with the appropriate antibiotics as follows: Blast (20 µg/ml) and puromycin (10 µg/ml). The organoid was passaged and seeded mixed with Matrigel for the next experiment.

### Subcutaneous organoid implantation

8-week-old male C57/B6J mice were obtained from the JAX and housed under specific pathogen-free conditions. The *APKS* tumoroids infected with sgNotum virus or sgControl were isolated from Matrigel using Cell Recovery Solution. The cell clusters (equivalent to 1 × 10^5^ cells for each mouse) were resuspended in 200 ul cold D-PBS with 0.5%BSA and 10% Matrigel and injected in the subcutaneous space of the loose skin between the shoulder blades. 4 weeks after transplantation, the tumor was isolated and weight.

### Western blot analysis

Organoids in 3D were digested with Cell Recovery Solution. The released organoids were washed with D-PBS and pelleted, then lysed with 1xRIPA Buffer (CST#9806) containing a protease/phosphatase inhibitor cocktail (CST #5872S). Organoids were disrupted by sonication and spun down for 15 min at 14,000 g. Supernatant was transferred to a clean tube and protein concentration determined by Bradford assay (Bio-Rad). An appropriate volume of 4x loading buffer was added to the cell lysates and incubated at 98°C for 10 min. Equal amounts of protein were loaded into each lane of an 8–12% SDS–PAGE gel. Membranes were blocked with 5% milk for 1 hour at room temperature and incubated with indicated primary antibodies diluted in proportion with 5% milk overnight at 4°C, followed by incubation with HRP conjugated secondary antibodies for 1 hour. Western blotting analyses were done using the following primary antibodies: anti-NOTUM (abcam#ab106448, 1:1,000), anti-β-ACTIN (abcam#ab6276, 1:5,000), anti-GPC1 (Proteintech#16700-1-AP, 1:1,000), anti-GPC4 (Proteintech#13048-1-AP, 1:1,000), anti-AKT (CST#4691, 1:1,000), anti-P-AKT (CST #4060, 1:1,000), anti-P38 (Proteintech #4064-1-AP, CST, 1:1,000), anti-phospho P38 (CST#4511, 1:1,000), anti-PS6(CST#4858, 1:1,000), anti-p-SMAD3(CST#9520, 1:1,000), anti-SMAD3(CST#9523, 1:1,000)and anti-SMAD4 (CST#46535, 1:1,000), anti-IIGFR1β(CST#3027S, 1:1,000), anti-TGFBR1(BOSTER#PB10101, 1:1,000). Signals were detected by HRP-conjugated secondary anti-rabbit antibody (CST#7074S, 1:2,000) and anti-mouse antibody (CST#7076S, 1:5,000) visualized with the SuperSignal West Pico PLUS Chemiluminescent Substrate (Thermo Scientific# 34577) and Bio-RAD Chemidoc TMMP imaging system.

### Co-immunoprecipitation assays

*Apc* mutant (*A*)and *Apc/Trp53* double mutant (*AP*) organoids were seeded with Matrigel and culture for 3 days. Treated the organoid with DMSO or ABC99(500 nm/ml) for 1 hour and then harvest the cells using 1× TrypLE. After washed with D-PBS, cell pellet was lysed using NP40 Cell Lysis Buffer (Thermo Fisher Scientific # FNN0021) containing a protease/phosphatase inhibitor cocktail (CST #5872S). Lysates were cleared by centrifugation at 12,000 rpm for 15 min. Transferred the supernatant to a new tube and discard the pellet containing cell debris and non-soluble fractions (save 50 ul supernatant for Input). Resuspended Dynabeads™ Protein A (Thermo Fisher Scientific # 10008D) and transferred 50 ul to a tube for each sample and incubated with 4 ug anti-GPC1 (Proteintech#16700-1-AP, 1:1,000) or anti-GPC4 (Proteintech#13048-1-AP, 1:1,000) or anti-Rabbit IgG (CST#3900) for 3 hours at room temperature. After washing using 200 ul PBS with Tween-20(0.01%), incubated the samples with beads overnight at 4°C on a rocking platform. Beads were then washed in lysis buffer and resuspended in 50 ul 2xSDS gel loading buffer and incubated at 70°C for 10 min. The proteins were separated on a 10% SDS-polyacrylamide gel, transferred to a PVDF membrane, and analyzed by immunoblotting with anti-IGF1Rβ (CST#3027S, 1:1,000) or anti-TGFBR1(BOSTER#PB10101, 1:1,000). Signals were detected by TidyBlot regent (BIO-RAD# STAR209PA) visualized with the SuperSignal West Pico PLUS Chemiluminescent Substrate (Thermo Scientific# 34577) and Bio-RAD Chemidoc TMMP imaging system.

### Clonal organoid formation assays

Organoids in 3D were digested into single cells using TrypLE™ Express Enzyme (1X) (Thermo Fisher#12605) for 5 min in 37°C water bath. Cell numbers were determined using a hemocytometer and single viable cells seeded in 24 well plates with Matrigel at the density of 500 cells/well. Each cell type was seeded in triplicate. Brightfield images of each well were captured 4 days post-seeding using microscope DMi8 (Leica, Wetzlar, Germany) and manually counted by using Photoshop software.

### Organoid proliferation assays

EdU assays: Organoids/tumoroids with varying genetic mutations infected with overexpression, knockdown, or control viruses were digested into single cells using TrypLE™ then seeded into 24-well plates at a density of 3,000 cells/well, with each cell type seeded in triplicate. Organoids were cultured for 4 days, with media changes every other day, after which 10 uM EdU (Thermo Fisher #E10187) was added to the culture media for 1-2 hours. EdU staining was performed with a Click-iT Plus EdU Alexa Fluor 647 Flow Cytometry Assay Kit (Thermo Fisher #10634), according to the manufacturer’s protocol. Stained cells were run on LSR Fortessa (BD Biosciences). The data were analyzed using Flowjo (BD Biosciences) software.

MTT assays: Organoids/tumoroids with varying genetic mutations infected with overexpression, knockdown, or control viruses were digested and seeded into 96-well plates at a density of 1,000 cells/well, each cell type seeded in 12 wells. Cell proliferation assays were performed using MTT (3-[4, 5-dimethylthiazol-2-yl]-2, 5-diphenyltetrazolium bromide) assays at indicated time points. Absorbance was measured at 560 nm. All data ware expressed as a mean from at least three independent biological experiments with n=3 per condition per experiment.

### Immunohistochemistry and Immunofluorescence

Tumors from mice were washed with cold DPBS, fixed in 4%PFA (Electron Microscopy Sciences# 15710), paraffin-embedded and sectioned. Haematoxylin & eosin, staining was performed in the Morphology Core of the UPenn NIDDK P30 Center for Molecular Studies in Digestive and Liver Diseases. For immunostaining, antigen retrieval was performed by heating slides in 0.01 M Tris-EDTA (pH 9.0) with a pressure cooker. For immunohistochemistry, the sections were then immunostained by the ABC peroxidase method (Vector Laboratories# PK-4001). For immunofluorescence staining, the sections were blocked with 10% donkey in PBS for 1 hour and incubated with primary antibodies at the appropriate dilution overnight in cold room. The next day incubated with Cy2- or Cy3-conjugated fluorescent secondary antibodies (Jackson Laboratory) and counterstained with DAPI in mounting media (Vector Laboratories). The following antibodies were used: Ki67 (Abcam#ab15580, 1:500), E-cadherin (CST#3195T,1:200), Notum (Abcam#ab106448, 1:500).

### Annexin V/PI Staining Assay

We seeded *APKS* organoid into 24 well plates and treated with ABC99 (500 nM/mL) or SB202190(10 uM/mL) or both for 2 days, using DMSO as vehicle control. For apoptosis detection, we used Dead Cell Apoptosis Kits (Thermo Scientific#V13245) and the analysis was performed according to the manufacturer’s instructions. The cell fluorescence was determined by flow cytometry, measuring the fluorescence emission at 530 nm and 575 nm (or equivalent) using 488 nm excitation.

### Single cell RNA sequencing

The normal colon was opened and cut longitudinally, incubated with 1U/ml dispase (STEMCELL Technologies# 07923) supplemented with 0.05 mg/ml liberase (Sigma-Aldrich# 05401127001) for 1 hours in 37°C water bath after mincing into small pieces. Tumors from *Apc^Min/+^* mice, AOM/DSS mice, or mice with tumors generated by orthotopic injection of *APKS* tumoroids were isolated and digested using dispase (1U/ml) with liberase (0.05 mg/ml). Tumor suspensions were shaken every ten minutes, then passed through 70 um cell strainers. Cells were washed with 0.04%BSA-DPBS and then stained with Cell Multiplexing Oligo (10X Genomics # 1000261) and incubated for 5 min (room temperature). After washing per the 10x Genomics protocol, cells were resuspended in 0.04%BSA-DPBS with DAPI and live single cells were FACS-purified. Library preparation was performed using the10x Chromium Next GEM Single Cell 3’ Reagent Kits v3.1 (Dual Index) according to manufacturer’s instructions (10x Genomics), approximately 2000-5000 cells were partitioned into Gel Beads in Emulsion (GEMs) with cell lysis and barcoded by reverse transcription of mRNA into cDNA, followed by amplification, enzymatic fragmentation and 5’ adaptor and sample index attachment. The recovery rate was approximately 800 cells per sample after filtering for quality control. Sample libraries were sequenced on the NextSeq 550 (Illumina).

### Quantitative RT-PCR

Colonoids, tumoroids, mouse, and human tissue samples were lysed using TRIzol (Invitrogen# 15596026) and total RNA was isolated. 1 µg total RNA was used in cDNA synthesis using RevertAid First Strand cDNA Synthesis Kit (Thermo Fisher Scientific#K1622) following manufacturer’s instruction. Quantitative PCR reactions were carried out with SYBR green (Applied Biosystems # 4367659) under standard conditions using QuantiStudio 7 Flex RealTime PCR (Applied Biosystems) and data were analyzed using QuantStudio RT-PCR software. Custom primers were validated with standard SYBR green qRT-PCR followed by melting curve analysis. Data were normalized to housekeeping gene *Gapdh*. The primers that were used to assess mouse *Notum*, human *NOTUM*, *Ascl2*, *Axin2*, *Ccnd1* and *Lgr5* mRNA expression levels were listed in **Supplementary table 3**.

### Bulk RNA sequencing

*Apc* mutant or *Apc/Trp53* double mutant tumoroids with or without Notum knockdown were harvested and lysed with TRIzol. After phase separation using the standard TRIzol protocol, the aqueous phase containing RNA was mixed with 70% ethanol and transferred to RNeasy MinElute spin columns (QIAGEN). Cleanup and further concentration of the RNA was done using the standard procedure of RNeasy MinElute Cleanup Kit.

CDNA synthesis was done using SMART-Seq v4 Ultra Low Input RNA Kit for Sequencing (Clontech) and the library was prepared using Nextera XT DNA Library Prep kit (Illumina) and sequenced using Illumina NextSeq 550. Sequencing of mRNA libraries generated 20-40 million high-quality 75-bp reads/sample. Adapters were trimmed using Trimmomatic v0.32 ^[47]^ and aligned to the mm10 genome using STAR v2.3.0e ^[48]^. Samtools v0.1.19 ^[49]^ was used to convert sam files to bam files, keep concordant primary alignments, sort and index. Rsubread’s ^[50]^ feature Counts command quantified reads per gene using Ensembl mm10 gene annotations. We removed all lowly expressed genes that did not have CPM >1 in at least a quarter of the samples. Limma v3.48.3^[51]^ voom was used to log2 transform and normalize the dataset. Limma was again used to fit a linear model and find differential genes. Camera^[52]^was used with EnsDb.Mmusculus.v75, as implemented with RNA-webInterface (https://github.com/bapoorva/RNA-WebInterface) to perform Gene Set Enrichment Analysis.

## Data Availability

The mouse transcriptome sequencing datasets generated during the current study will be made available in the NCBI Gene Expression Omnibus (GEO). Human single cell RNA sequencing data analyzed were collected under an active IRB and the study describing their collection is currently in preparation. Human scRNAseq data is being deposited at dbGaP. Other software tools (including version numbers) for RNA-seq, and scRNA-seq analyses are listed in the Key resources table.

## Extended tables

**Supplementary table1:**
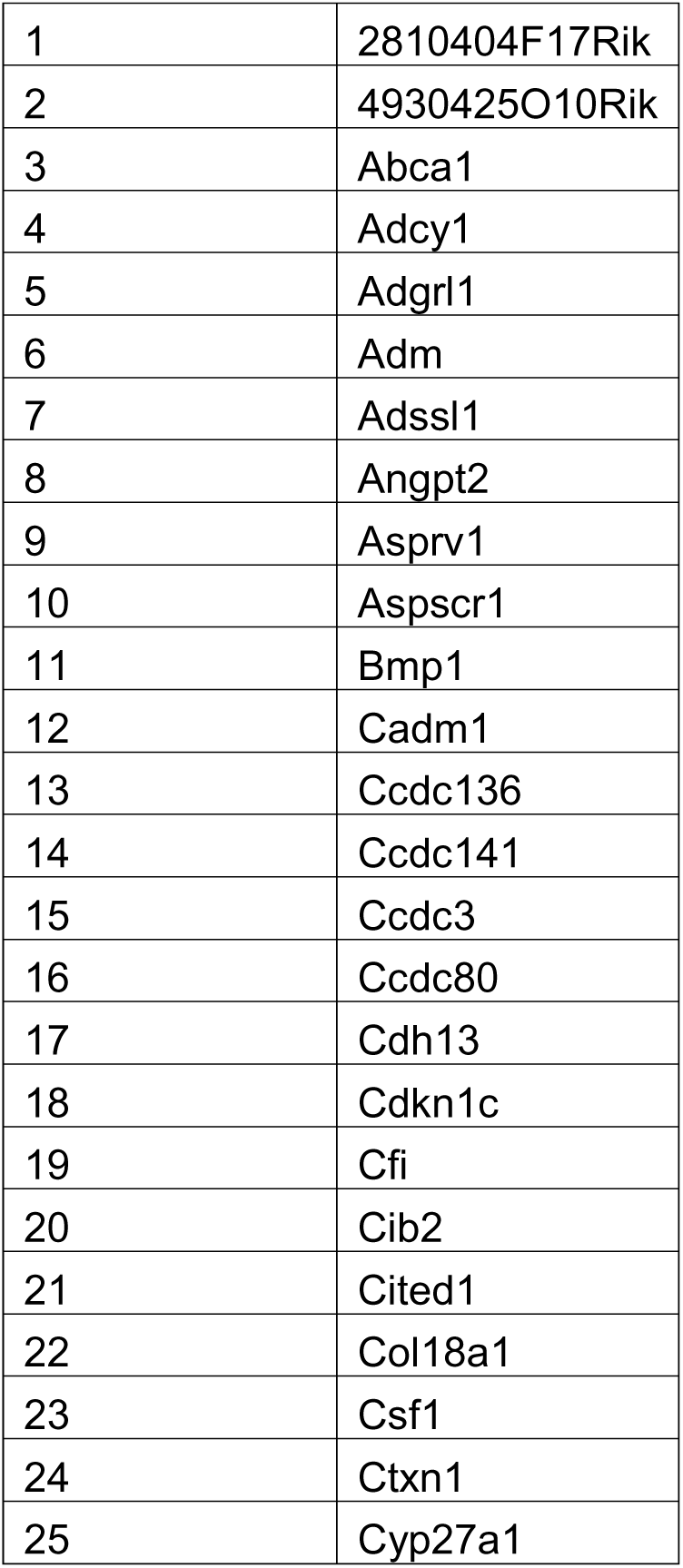
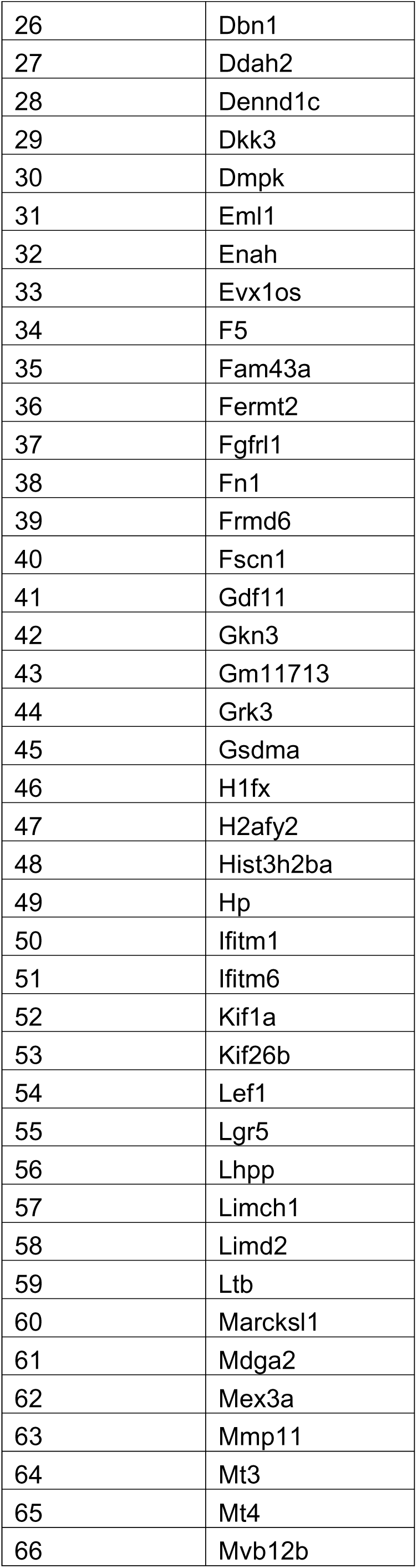
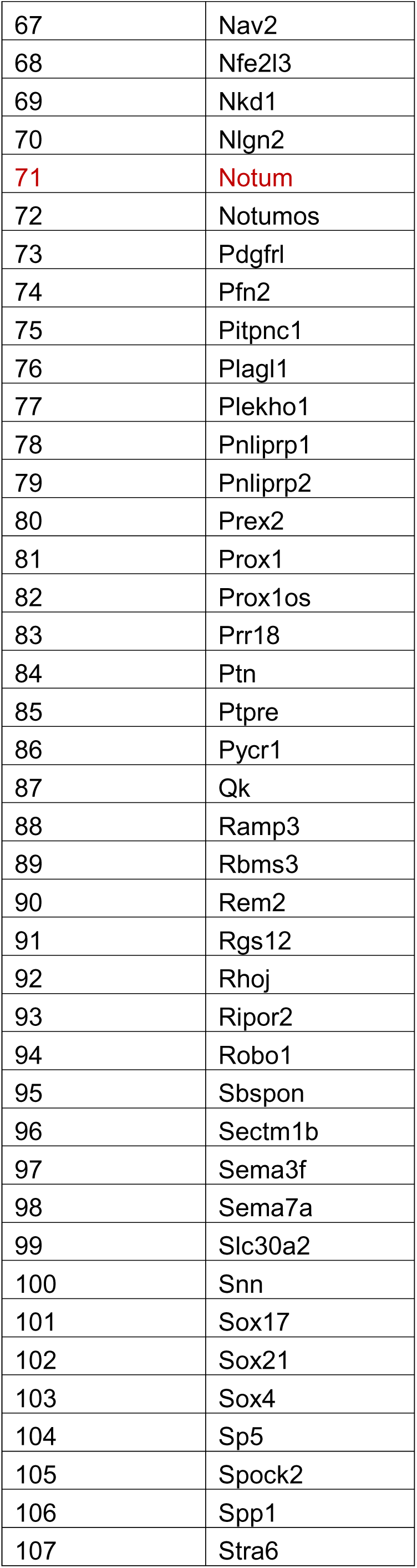
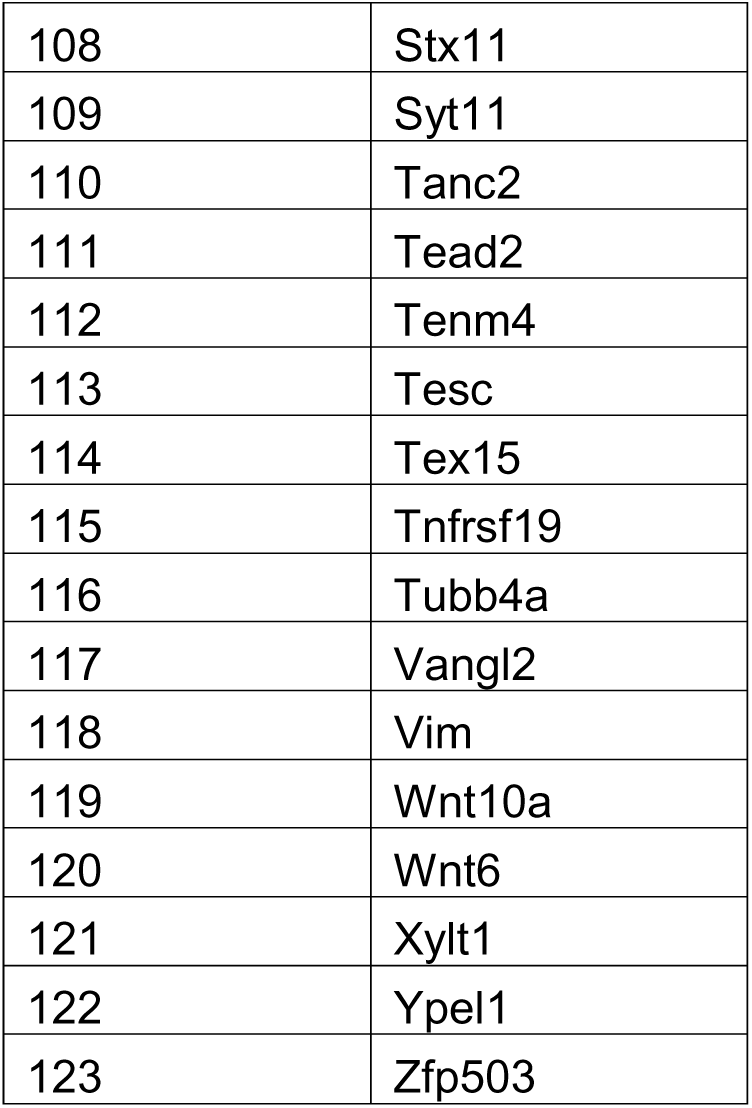
genes that acutely induced upon APC loss and remains elevated in colon cancer models.

**Supplementary table 2:** Transcriptional pathways most differentially affected by NOTUM LOF between ApcΔ and AP double mutant cultures.

| A ctrl vs A Notum shRNA |  | Up= Up in Notum shRNA |  |  |
| --- | --- | --- | --- | --- |
| name | NGenes | Direction | p-Value | FDR |
| HALLMARK_E2F_TARGETS | 202 | Up | 5.728E-05 | 0.00281891 |
| HALLMARK_CHOLESTEROL_HOMEOSTASIS | 70 | Up | 0.00016627 | 0.00281891 |
| HALLMARK_PROTEIN_SECRETION | 92 | Down | 0.00017475 | 0.00281891 |
| HALLMARK_G2M_CHECKPOINT | 199 | Up | 0.00022551 | 0.00281891 |
| HALLMARK_ALLOGRAFT_REJECTION | 116 | Up | 0.00709347 | 0.07093473 |
| HALLMARK_MTORC1_SIGNALING | 193 | Up | 0.01053037 | 0.08775306 |
| HALLMARK_OXIDATIVE_PHOSPHORYLATION | 192 | Down | 0.01637338 | 0.11695273 |
| HALLMARK_INTERFERON_ALPHA_RESPONSE | 90 | Up | 0.01942423 | 0.12000662 |
| HALLMARK_P53_PATHWAY | 183 | Down | 0.02177959 | 0.12000662 |
| HALLMARK_INTERFERON_GAMMA_RESPONSE | 160 | Up | 0.02400132 | 0.12000662 |
| AP ctrl vs AP Notum shRNA |  | Up= Up in Notum shRNA |  |  |
| name | NGenes | Direction | p-Value | FDR |
| HALLMARK_INTERFERON_ALPHA_RESPONSE | 90 | Down | 3.10E-09 | 1.55E-07 |
| HALLMARK_TNFA_SIGNALING_VIA_NFKB | 177 | Up | 5.7907E-06 | 0.00014477 |
| HALLMARK_FATTY_ACID_METABOLISM | 135 | Down | 0.00055716 | 0.009139 |
| HALLMARK_MYC_TARGETS_V2 | 58 | Up | 0.00073112 | 0.009139 |
| HALLMARK_INTERFERON_GAMMA_RESPONSE | 160 | Down | 0.00132664 | 0.01326643 |
| HALLMARK_UNFOLDED_PROTEIN_RESPONSE | 110 | Up | 0.00421123 | 0.0350936 |
| HALLMARK_TGF_BETA_SIGNALING | 50 | Up | 0.00559056 | 0.03993258 |
| HALLMARK_MYC_TARGETS_V1 | 202 | Up | 0.00912038 | 0.05700236 |
| HALLMARK_G2M_CHECKPOINT | 199 | Up | 0.01108253 | 0.06156962 |
| HALLMARK_ADIPOGENESIS | 183 | Down | 0.01708474 | 0.08542368 |

**Supplementary table 3:**
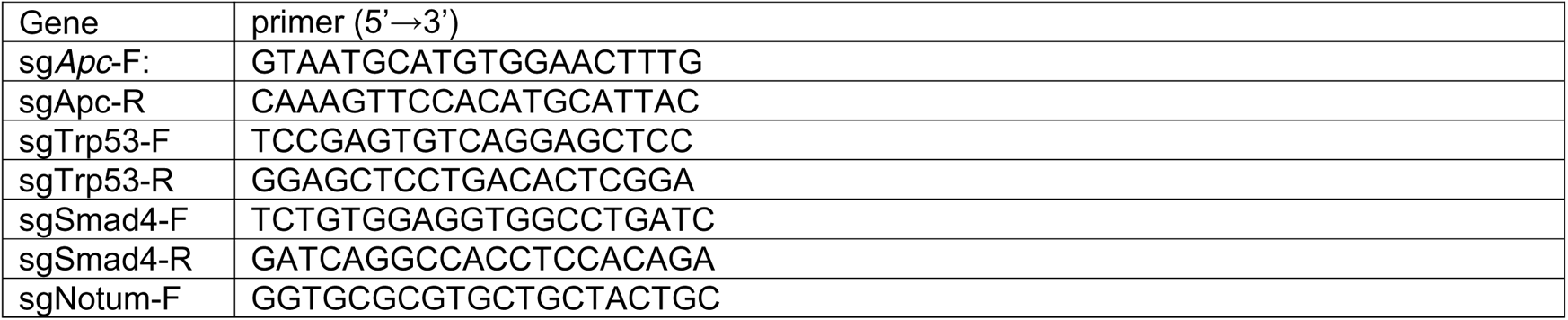
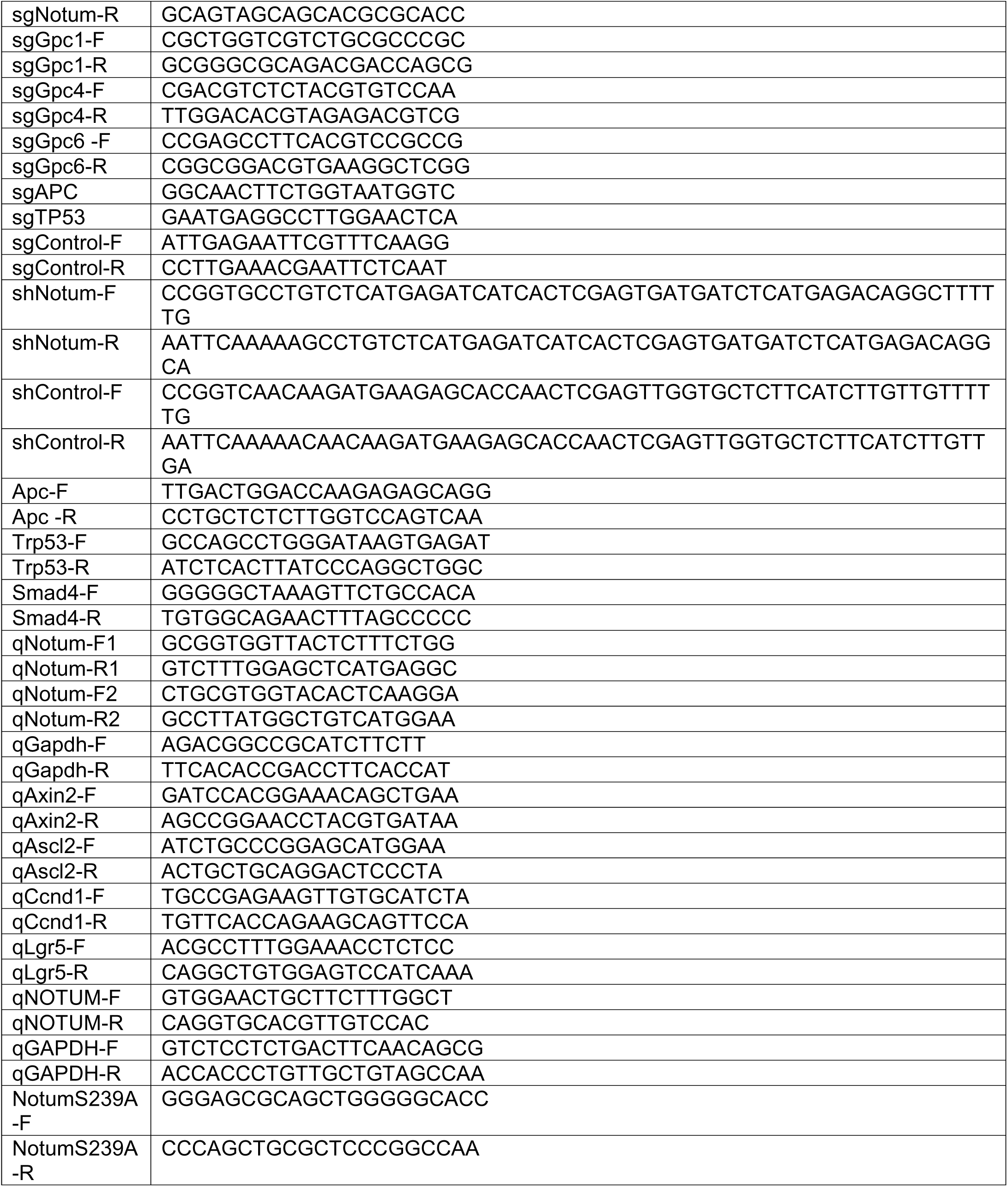
Primers that used in this paper.

---

**Extended Data Fig. 1.**
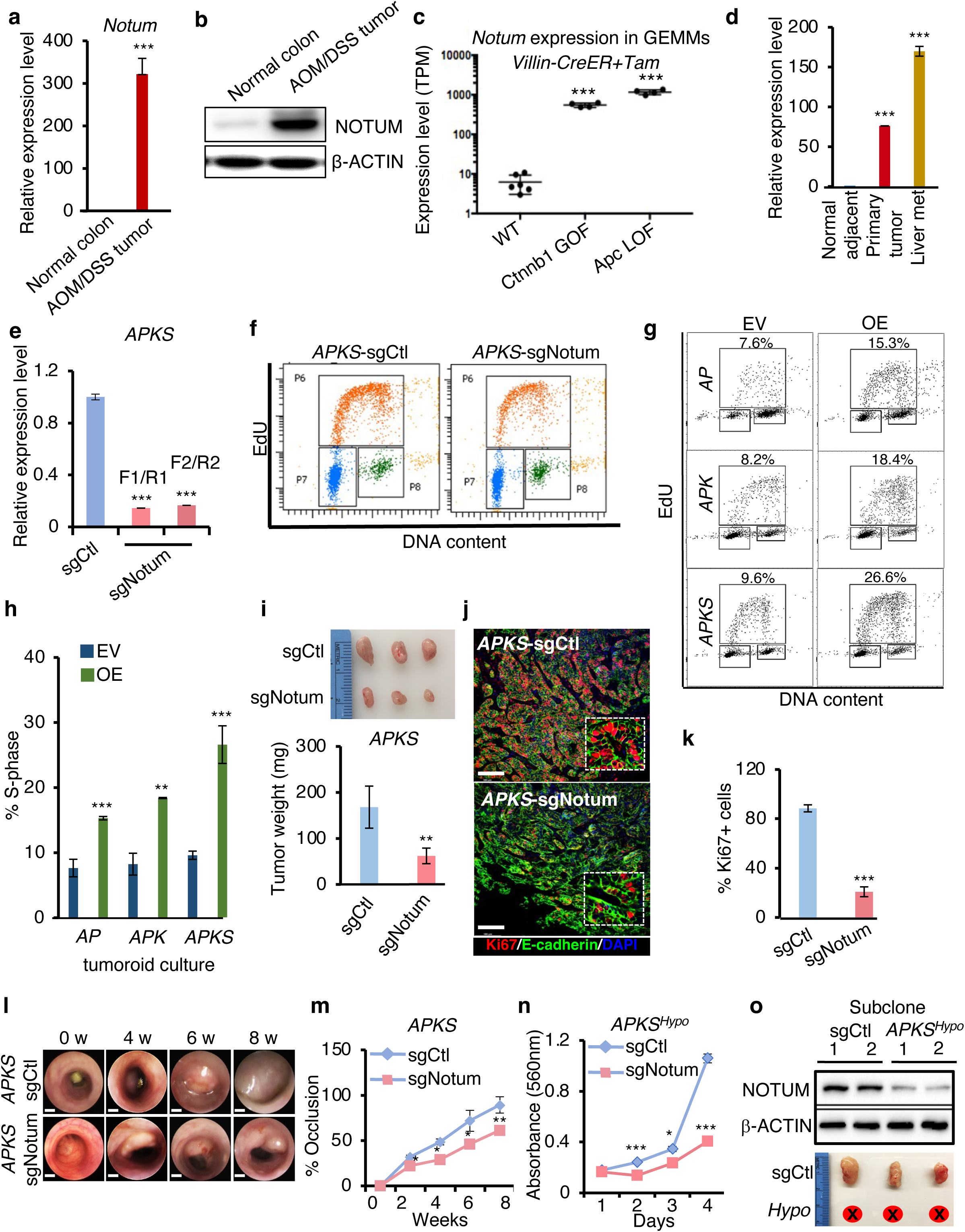
*Notum* is an obligate oncogene in advanced adenocarcinoma models. **(a)** Quantitative RT-PCR analysis of *Notum* expression in normal colon epithelial cells and AOM/DSS carcinoma cells (n=3). **(b)** Western blot analysis for NOTUM in normal colon epithelial cells and AOM/DSS carcinoma cells. β-ACTIN was used as a loading control. **(c)** Expression of *Notum* in colonic epithelium of *Ctnnb1^exon3flox/flox^::Villin-CreER* and *Apc^flox/flox^::VillinCreER* mice 5 days after tamoxifen-induced gene ablation, relative to controls (n=6). **(d)** Quantitative RT-PCR for *NOTUM* in human primary colorectal adenocarcinoma, cognate liver metastasis, and normal adjacent tissue colon (n=6). **(e)** Expression of *Notum* in *APKS* tumoroids infected with control sgRNA (sgCtl) or *Notum* sgRNA (sgNotum) (n=3). **(f)** Flow cytometric analysis of EdU incorporation in *APKS* tumoroids infected with control sgRNA (sgCtl) or *Notum* sgRNA (sgNotum), quantified in **Fig.1h** (n=3). **(g)** Flow cytometric analysis of EdU incorporation in *AP*, *APK* and *APKS* after infection with empty lentiviral vector (pUltra-EV) or vector overexpressing NOTUM (Notum OE) (n=3). **(h)** Percentage of S phase cells from (**g**). **(i)** Top: Subcutaneous tumors formed 4 weeks after implantation of *APKS* tumoroids infected with control sgRNA (sgCtl) or *Notum* sgRNA (sgNotum) (n=3). Bottom: Weight of tumors above. **(j)** Immunofluorescence staining for KI67 and E-CADHERIN on tumors formed from subcutaneous *APKS* injection. Nuclei are visualized by DAPI staining (n=3). Scale bar: 100 *μ*m. **(k)** Quantification of KI67+ cells in subcutaneous *APKS-*sgCtl and *APKS-*sgNotum tumors. **(l)** Representative endoscopic pictures showing the development of orthotopically implanted *APKS* tumors with control sgRNA (sgCtl) or *Notum* sgRNA (sgNotum) over 8 weeks of tumor growth (n=5 in sgCtl group, n=4 in sgNotum group). Scale bar: 2 mm. **(m)** The percent of lumen occlusion was measured for each *APKS* tumor from (**l**) during the course of the experiment. **(n)** MTT assays in a subclone of *APKS* tumoroids infected with *Notum* sgRNA (sgNotum) and exhibiting hypomorphic Notum loss of function (*APKS^Hypo^*) (n=3). **(o)** Top: Western blot analysis of NOTUN expression in *APKS^Hypo^* subclone from (N). β-ACTIN was used as a loading control. Bottom: Subcutaneous tumor formation using the Notum *APKS^Hypo^* line or control APKS tumoroids. For all panels: \**P*<0.05, \*\**P*<0.01, \*\*\**P*<0.001, Student’s *t*-test.

**Extended Data Fig.2.**
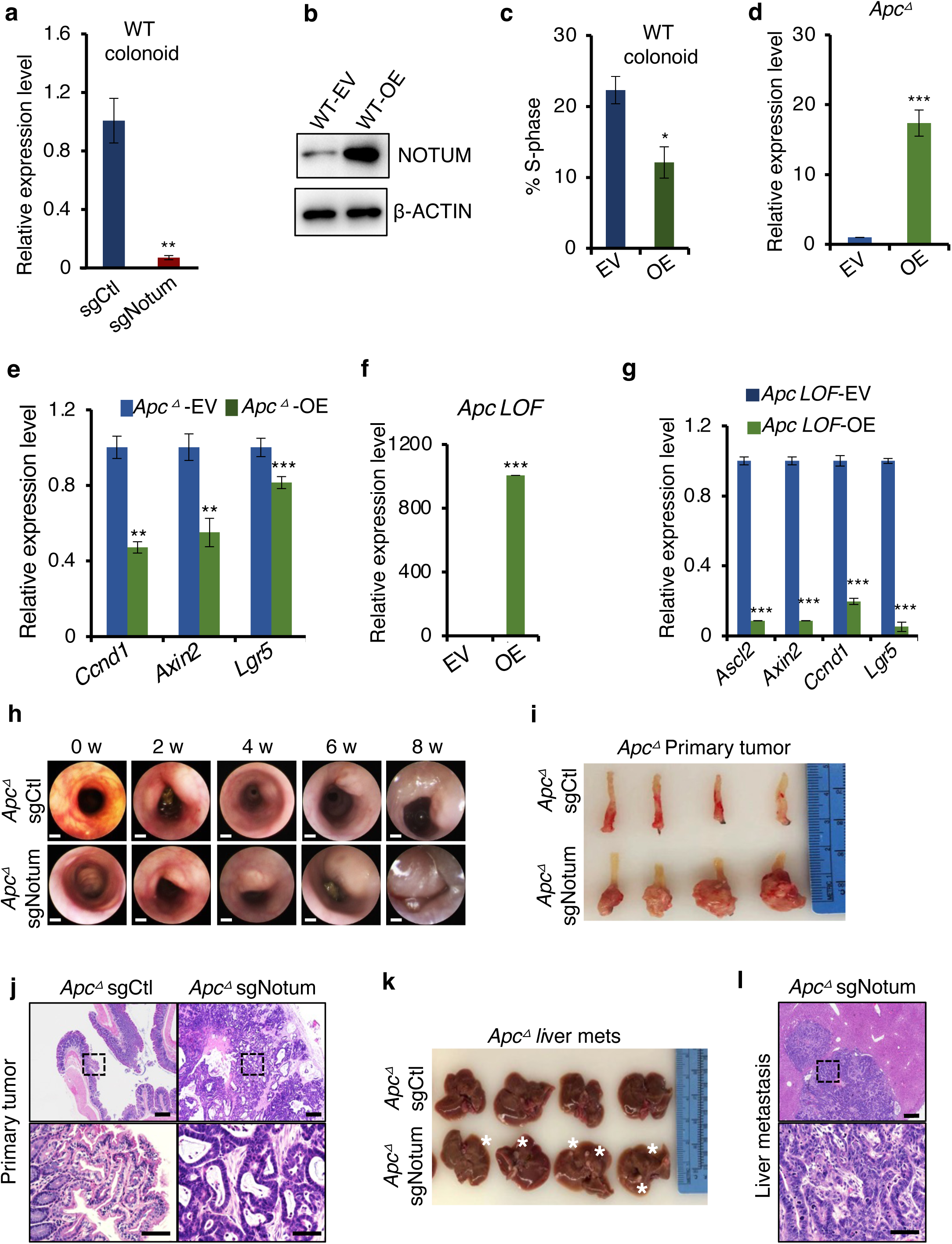
NOTUM maintains tumor suppressive activity upon APC inactivation. **(a)** Expression level of *Notum* in wildtype organoids infected with control sgRNA (sgCtl) or *Notum* sgRNA (sgNotum) (n=3). **(b)** Western blot analysis for NOTUM in wildtype organoids infected with empty vector (EV) or *Notum* overexpression plasmid (OE). β-ACTIN was used as a loading control. **(c)** The percentage of EdU+ cells in wildtype organoids infected with empty vector (EV) or *Notum* overexpression plasmid (OE) (n=3). **(d)** Expression level of *Notum* in *Apc* mutant tumoroids infected with empty vector (EV) or vector overexpressing *Notum* (OE) (n=3). **(e)** Quantitative RT-PCR for canonical Wnt/ β-CATENIN pathway target genes in *Apc* mutant tumoroids from (**d**). **(f)** Expression level of *Notum* in *Apc^flox/flox^::Villin-CreER* colon tumoroids derived after tamoxifen treatment, infected with empty vector (EV) or vector overexpressing *Notum* (OE) (n=3). **(g)** Quantitative RT-PCR for canonical Wnt/ β-CATENIN pathway target genes in *Apc* mutant tumoroids from *Apc^flox/flox^::Villin-CreER* mice from (**f**). **(h)** Representative endoscopic pictures showing the development of orthotopic tumors formed in the distal colon by *Apc* mutant tumoroids infected with control sgRNA (sgCtl) or *Notum* sgRNA (sgNotum) (n=4). Scale bar: 2 mm. **(i)** Orthotopic primary tumors from (**h**). **(j)** Hematoxylin and eosin micrographs from tumors in (**i**). Scale bar of top panels: 200 *μ*m. lower panels: 50 *μ*m. **(k)** Livers of mice from (**h**), With white asterisks indicating macroscopically visible metastatic lesions. **(l)** Hematoxylin and eosin micrographs from liver metastases in (**k**). Scale bar of top panel: 200 *μ*m. Lower panel: 50 *μ*m. For all panels: \**P*<0.05, \*\**P*<0.01, \*\*\**P*<0.001, Student’s *t*-test.

**Extended Data Fig.3.**
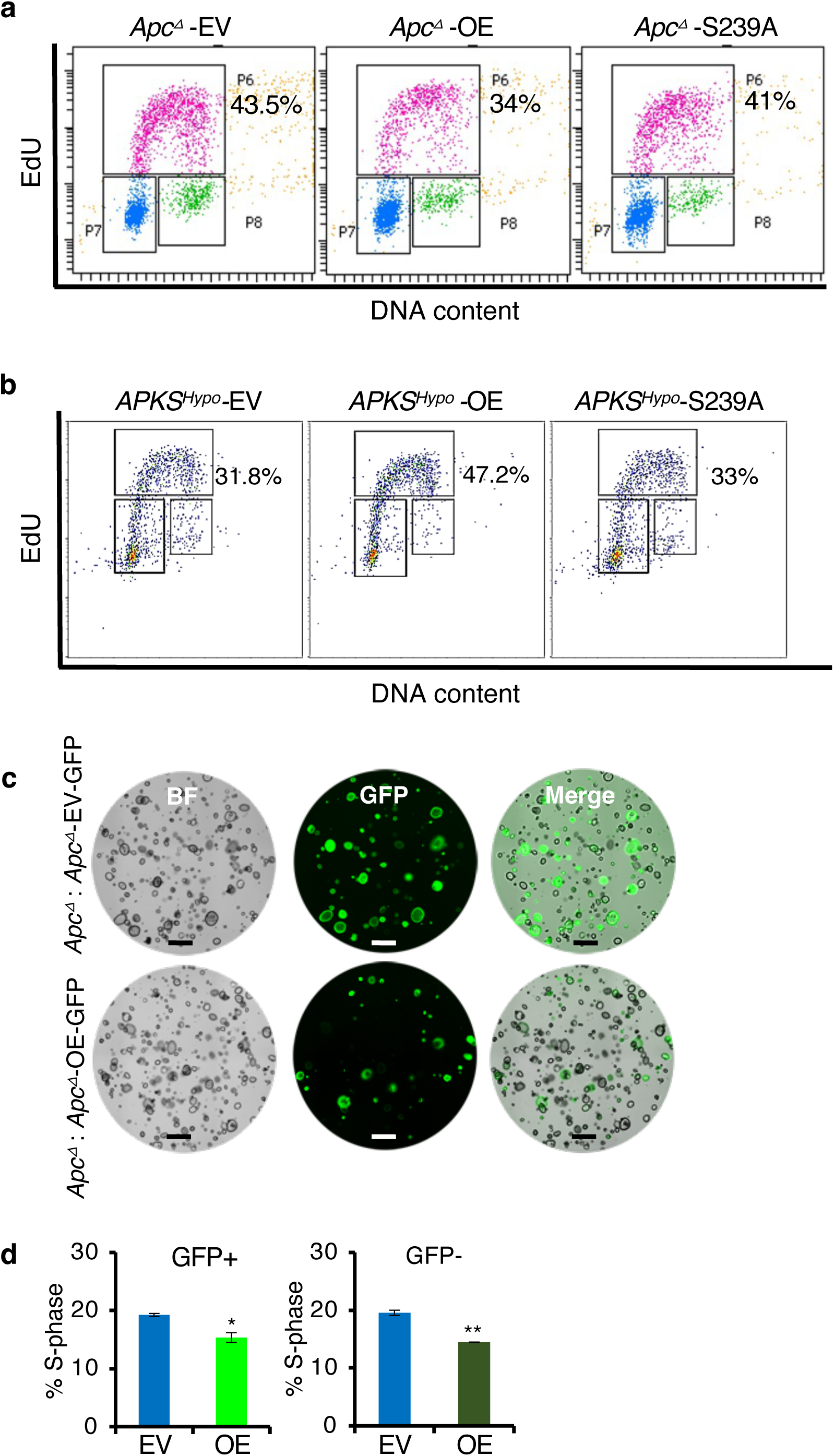
Novel tumor suppressive and oncogenic functions of NOTUM require extracellular enzymatic activity. **(a)** Representative flow cytometric plots of EdU incorporation after a 2-hour pulse, from Fig. 3a, c: *Apc* mutant tumoroids infected with empty vector (EV), vector expressing wildtype NOTUM (OE), or catalytically dead NOTUM with Ser239Ala mutation (S239A) (n=3). **(b)** Representative flow cytometric plots of EdU incorporation after a 2-hour pulse, from Fig. 3d, f: *APKS^Hypo^* tumoroids infected with empty vector (EV), vector expressing wildtype NOTUM (OE), or catalytically dead NOTUM with Ser239Ala mutation (S239A) (n=3). **(c)** Images of *Apc*-mutant tumoroid co-cultures, with half of the tumoroids infected with either a vector expressing GFP only (EV-GFP) or expressing GFP and NOTUM (OE-GFP), and the other half uninfected (n=3). Scale bar: 100 *μ*m. **(d)** EdU assays in the cultures in (c), quantifying the percentage of cells in S phase in the GFP+ and GFP-populations. \**P*<0.05, \*\**P*<0.01, Student’s *t*-test.

**Extended Data Fig.4.**
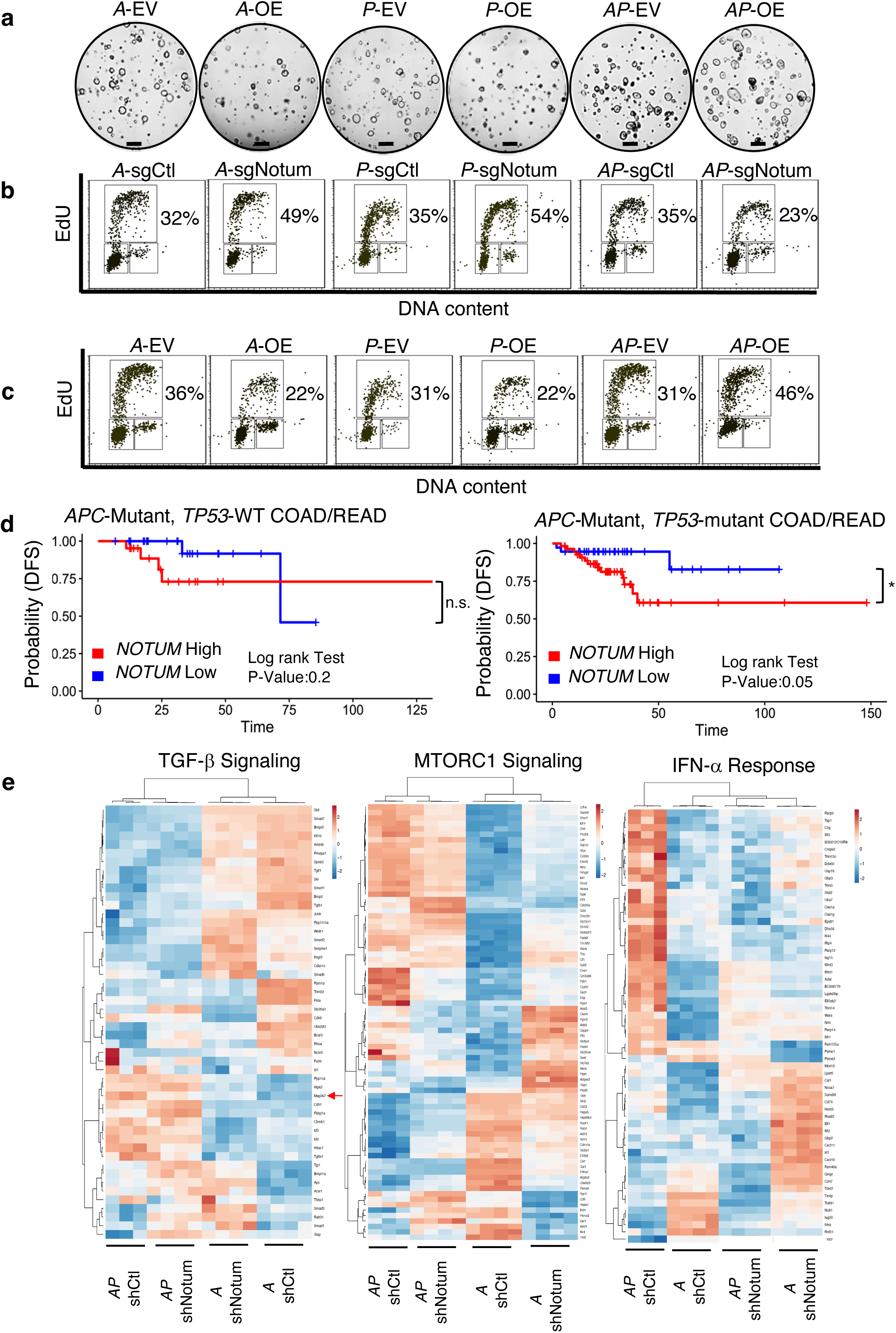
*Apc* and *Trp53* inactivation synergize to confer oncogenic activity upon NOTUM. **(a)** Brightfield image of *A*, *P* and *AP* tumoroids infected with vector overexpressing NOTUM (OE) or empty vector control (EV) (n=3). Scale bar: 100 *μ*m. **(b)** Representative flow cytometric plots of EdU incorporation after a 2-hour pulse, from Fig. 4b, d: *Apc* (*A*), *Trp53* (*P*), or double mutant (*AP*) tumoroids infected with control sgRNA (sgCtl) or *Notum* sgRNA (sgNotum) (n=3). **(c)** Representative flow cytometric plots of EdU incorporation after a 2-hour pulse, from Fig. 4f, Extended Data Fig 4a: *A*, *P* and *AP* tumoroids infected with vector overexpressing NOTUM (OE) or empty vector control (EV) (n=3). **(d)** Analysis of disease-free survival (DFS) in colon and rectal adenocarcinoma patients (COAD, READ) for which gene expression data is available in the Cancer Genome Atlas (TCGA). Left panel shows DFS for *APC* mutant, *TP53* wildtype patients (n=46), binned on *NOTUM* expression above (high) or below (low) the median. Right panel shows DFS for *APC/TP53* double mutant COAD/READ patients (n=93), binned on *NOTUM* expression above (high) or below (low) the median). \**P*<0.05, Student’s t-test. **(e)** Heatmaps showing gene expression in gene sets identified by gene set enrichment analysis for pathways affected by NOTUM loss of function in *A* or *AP* tumoroids.

**Extended Data Fig.5.**
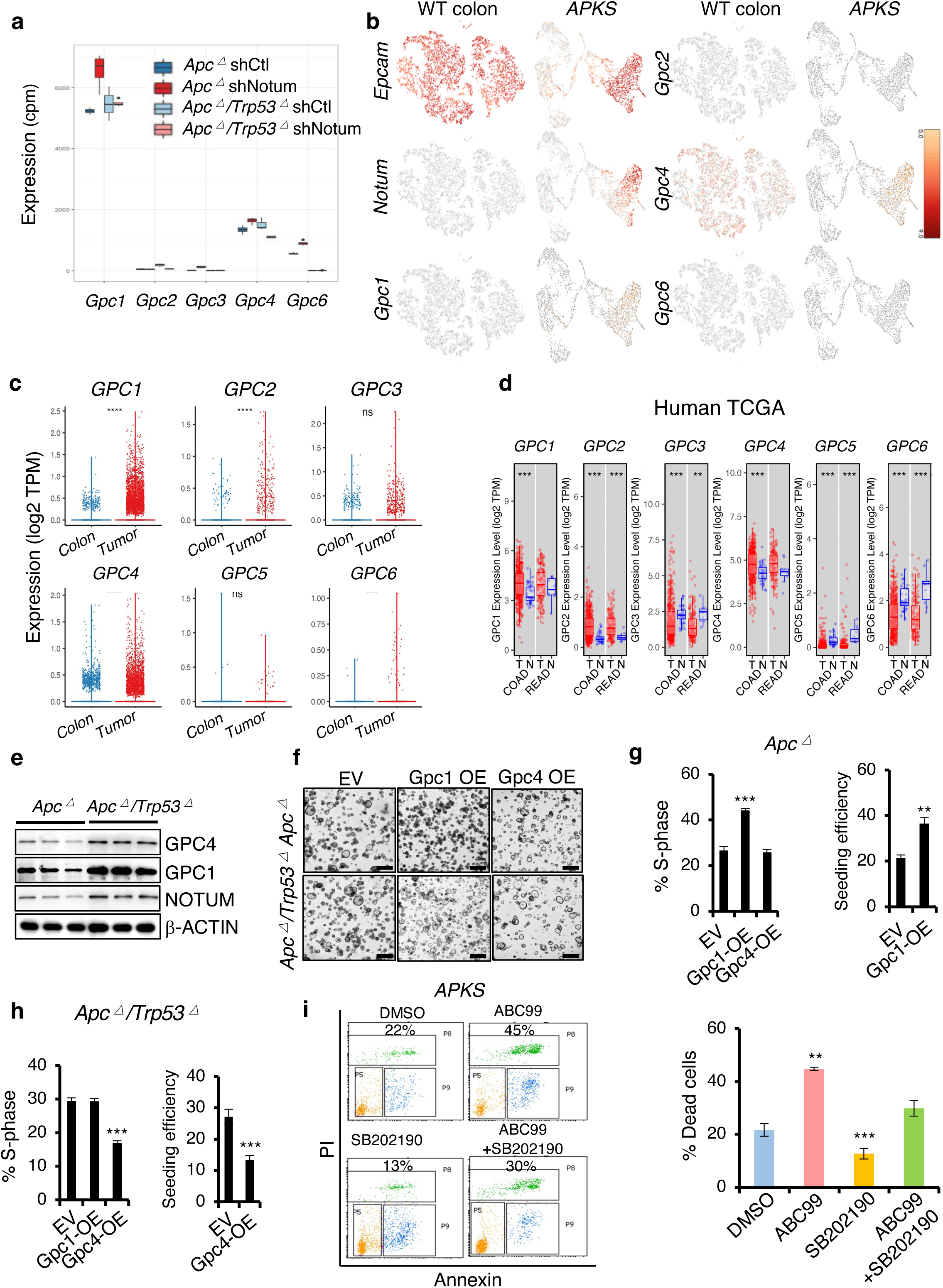
Glypicans as mediators of NOTUM function in adenoma and adenocarcinoma. **(a)** Expression level of Glypicans *Gpc1, Gpc2, Gpc3, Gpc4* and *Gpc6* (*Gpc5* was not detected) in RNA sequencing data from *Apc* mutant (*A*) and *Apc/Trp53* double mutant (*AP*) tumoroids infected with control shRNA (shCtl) or *Notum* shRNA (shNotum) (n=4). **(b)** UMAP plots showing expression of *Epcam, Notum, Gpc1, Gpc2, Gpc4* and *Gpc6* in single cell RNA sequencing data from wildtype colon epithelium and adenocarcinoma derived from orthotopic implantation of *APKS* tumoroids into syngeneic mouse colonic mucosa. **(c)** Expression level of Glypicans *GPC1, GPC2, GPC3, GPC4, GPC5*, and *GPC6* in single cell RNAseq data from primary human colorectal cancers and normal adjacent colon (n=17). **(d)** Expression level of *GPC1, GPC2, GPC3, GPC4, GPC5* and *GPC6* in human TCGA colon (COAD) and rectum (READ) adenocarcinoma datasets. **(e)** Western blotting for GPC1 and 4 in whole cell lysates from *A* and *AP* tumoroids. β-ACTIN was used as a loading control. **(f)** Brightfield image of *A* and *AP* tumoroids infected with empty vector (EV) or vector expressing GPC1 (Gpc1 OE) or GPC4 (Gpc4 OE) (n=3). Scale bar: 100 *μ*m. **(g,h)** EdU assays (left) and clonal seeding efficiencies (right) from *Apc* mutant (**g**) and *Apc/Trp53* double mutant (**h**) cultures shown in (**f**) (n=3). **(i)** Quantification of apoptosis in *APKS* tumoroid cultures after inhibition of NOTUM with the small molecule ABC99, the p38 inhibitor SB202190, or both (n=3). For all panels: ****P<0.0001, \*\*\**P*<0.001, \*\**P*<0.01, Student’s *t*-test.

**Extended Data Fig.6.**
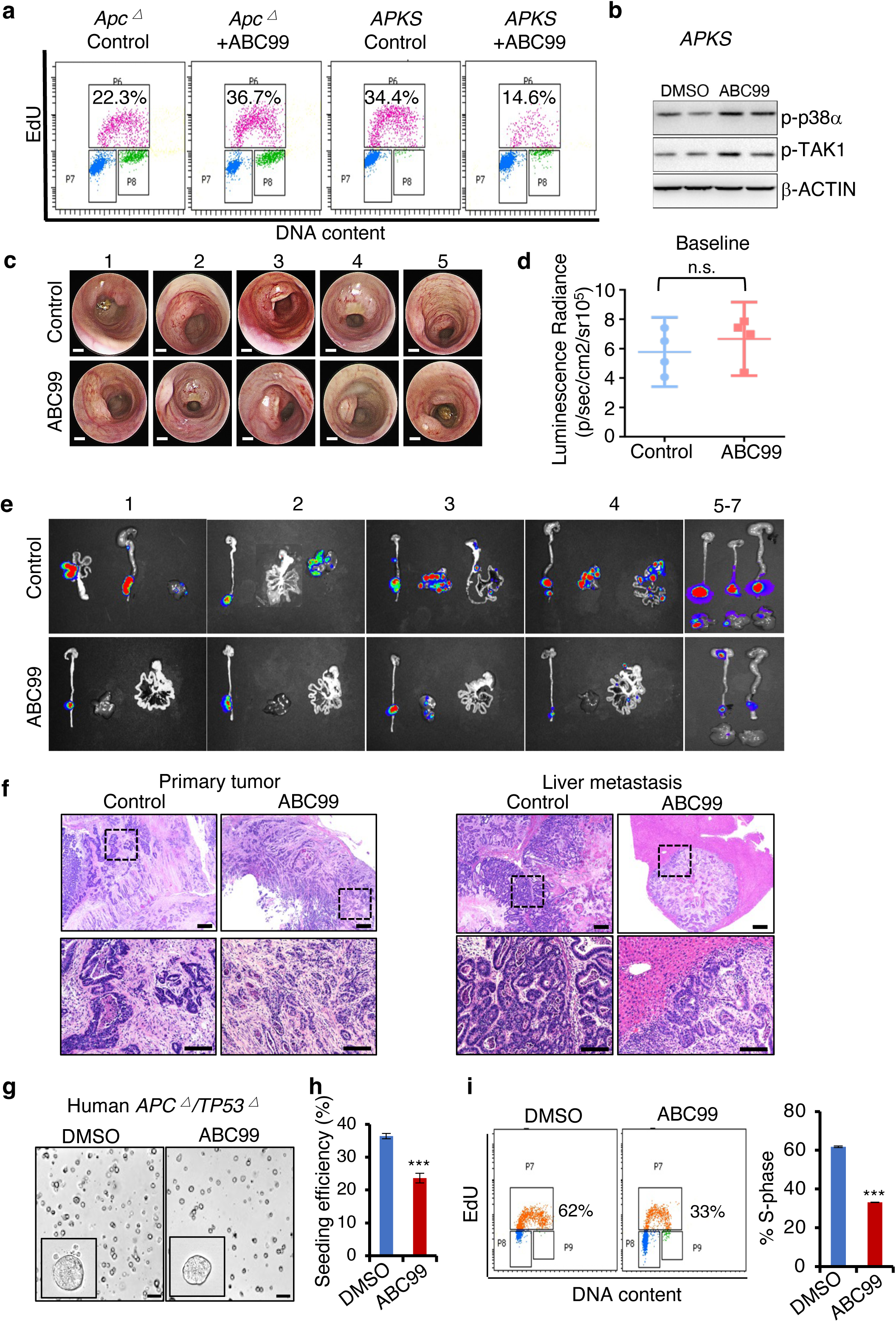
Pharmacological NOTUM inhibition is efficacious in a pre-clinical animal model of metastatic colon cancer. **(a)** Representative flow cytometric analysis of EdU incorporation in *Apc* mutant and APKS tumoroids treated with the small molecule inhibitor of NOTUM, ABC99, or vehicle control (DMSO), quantified in **Fig. 6a** (n=3). **(b)** Western blotting for phosphorylation of TAK1 and p38α in APKS tumoroids treated with vehicle or ABC99. β-ACTIN was used as a loading control. **(c)** Representative endoscopic images of the orthotopic APKS tumors from Fig. 6b prior to initiation of treatment, 4 weeks post-implantation. Scale bar: 2 mm. **(d)** Quantification of luminescence radiance from tumors in (**c**) after random assignment to groups and prior to initiation of treatment. **(e)** Bioluminescence images of dissected intestine, colon, and livers of representative mice from **Fig. 6c** at experimental endpoint. **(f)** Hematoxylin and Eosin-stained histological sections of representative primary tumors and liver metastases mice from **Fig. 6c** at the experimental endpoint. Scale bar of top panels: 200 *μ*m. Lower panels: 100 *μ*m. **(g)** Brightfield micrographs of human colon cultures harboring loss-of-function mutations in *APC* and *TP53*, treated with ABC99 or vehicle control (n=3). Scale bar: 100 *μ*m. **(h)** Clonal seeding efficiency from single cells from cultures shown in (**g**). **(i)** EdU incorporation assays in cultures shown in (**g**), quantified at right (n=3). For all panels: \*\*\**P*<0.001, Student’s *t*-test.

